# Disentangling signal and noise in neural responses through generative modeling

**DOI:** 10.1101/2024.04.22.590510

**Authors:** Kendrick Kay, Jacob S. Prince, Thomas Gebhart, Greta Tuckute, Jingyang Zhou, Thomas Naselaris, Heiko Schutt

**Affiliations:** Center for Magnetic Resonance Research (CMRR), Department of Radiology, University of Minnesota; Department of Psychology, Harvard University; Department of Computer Science, University of Minnesota; Department of Brain and Cognitive Sciences, Massachusetts Institute of Technology; McGovern Institute for Brain Research, Massachusetts Institute of Technology; Center for Computational Neuroscience (CCN), Flatiron Institute; Department of Neuroscience, University of Minnesota; Department of Behavioural and Cognitive Sciences, Université du Luxembourg

## Abstract

Measurements of neural responses to identically repeated experimental events often exhibit large amounts of variability. This *noise* is distinct from *signal*, operationally defined as the average expected response across repeated trials for each given event. Accurately distinguishing signal from noise is important, as each is a target that is worthy of study (many believe noise reflects important aspects of brain function) and it is important not to confuse one for the other. Here, we describe a principled modeling approach in which response measurements are explicitly modeled as the sum of samples from multivariate signal and noise distributions. In our proposed method—termed Generative Modeling of Signal and Noise (GSN)—the signal distribution is estimated by subtracting the estimated noise distribution from the estimated data distribution. Importantly, GSN improves estimates of the signal distribution, but does not provide improved estimates of responses to individual events. We validate GSN using ground-truth simulations and show that it compares favorably with related methods. We also demonstrate the application of GSN to empirical fMRI data to illustrate a simple consequence of GSN: by disentangling signal and noise components in neural responses, GSN denoises principal components analysis and improves estimates of dimensionality. We end by discussing other situations that may benefit from GSN’s characterization of signal and noise, such as estimation of noise ceilings for computational models of neural activity. A code toolbox for GSN is provided with both MATLAB and Python implementations.

## Introduction

Nominally identical sensory, cognitive, and/or motor events often result in highly variable neural activity measurements (Goris et al., 2014; Ito et al., 2020; Rabinowitz et al., 2015; Tolhurst et al., 1981). Such variability is termed *noise*, and manifests in all techniques for measuring brain activity, including electrophysiology, optical imaging, electroencephalography, magnetoencephalography, and functional magnetic resonance imaging (fMRI). Noise may originate from multiple sources. Noise can arise for instrumental reasons (e.g., electrical noise, head motion) or physiological reasons (e.g., cardiac noise), but can also reflect genuine variability in neural activity. Another important aspect of noise is its complex multivariate nature: variability in activity is not independent across units (e.g., neurons, voxels, channels) but typically exhibits structured correlations (Biswal et al., 1995; Cohen and Kohn, 2011; Hazon et al., 2022; Kanitscheider et al., 2015; Mell et al., 2021; Moreno-Bote et al., 2014). To mitigate the effects of noise, neuroscientists usually average neural responses across repeated trials associated with the same event. The underlying premise is that the object of interest, the *signal*, is not the neural response observed on any single trial but the average expected neural response across a large (infinite) number of trials.

Many research programs in systems, cognitive, and computational neuroscience focus on studying signal. For example, one might seek to characterize the tuning of sensory neurons by averaging responses across several trials measured for each stimulus condition. But there are also scientific motivations for characterizing and understanding noise, which may play an important role in neural computation (Panzeri et al., 2022; Ringach, 2009; Uddin, 2020). One example approach, originating in computational neuroscience, investigates the correlational structure of noise in the responses of individual neurons and explores how these noise correlations affect the information capacity of a neural population code (Averbeck et al., 2006; Azeredo da Silveira and Rieke, 2021; Cafaro and Rieke, 2010; Zylberberg et al., 2016). Another approach, commonly referred to as resting-state functional connectivity, leverages spontaneous activity fluctuations to parcellate brain areas and networks (Eickhoff et al., 2018) and to develop biomarkers for individuals (Gratton et al., 2020) or populations (Zhang et al., 2021). Perhaps the deepest potential interpretation of noise is that it reflects critical latent cognitive processes that are not directly controlled by the experimental paradigm. One example of this view is the theory that noise reflects statistical priors and/or probabilistic neural computations (Ma et al., 2006; Orbán et al., 2016; van Bergen et al., 2015).

Given that both signal and noise are of potential interest, a challenge faced by neuroscientists is that signal and noise are entangled in neural activity measurements, and it is not immediately obvious how to separate the two components. The standard approach is to average responses across trials and assume that the result adequately isolates signal from noise. However, while simple and straightforward, the approach of trial averaging does not necessarily produce perfectly accurate signal measures, a point that has been previously recognized (Pospisil and Bair, 2021a; Pospisil and Pillow, 2024; Stringer et al., 2019). To illustrate, we perform a simple simulation in which two units exhibit positive noise correlation but no signal correlation (**Figure 1**). When the number of trials per condition is large, trial averaging indeed suppresses the noise, but noise correlation is still observed in the trial-averaged results (panel A). When the number of trials per condition is small, noise correlation in the trial-averaged results is even more substantial (panel B). Finally, to further accentuate the point, we simulate a situation where there is no signal at all (panel C): this case clearly shows how noise structure seeps into the trial-averaged results. The residual influence of noise on trial-averaged results is a problem as it may lead to inaccurate estimates of signal correlation (Pospisil and Bair, 2021a), and inaccurate interpretations of commonly performed multivariate analyses, such as principal components analysis, representational similarity analysis, and analysis of neural dimensionality. In short, what is thought to be due to signal might actually be due to noise. Indeed, there has been recent interest in methods for identifying and isolating signal and noise components in high-dimensional neural data (Pospisil and Pillow, 2024; Stringer et al., 2019; Williams and Linderman, 2021).

**Figure 1.**
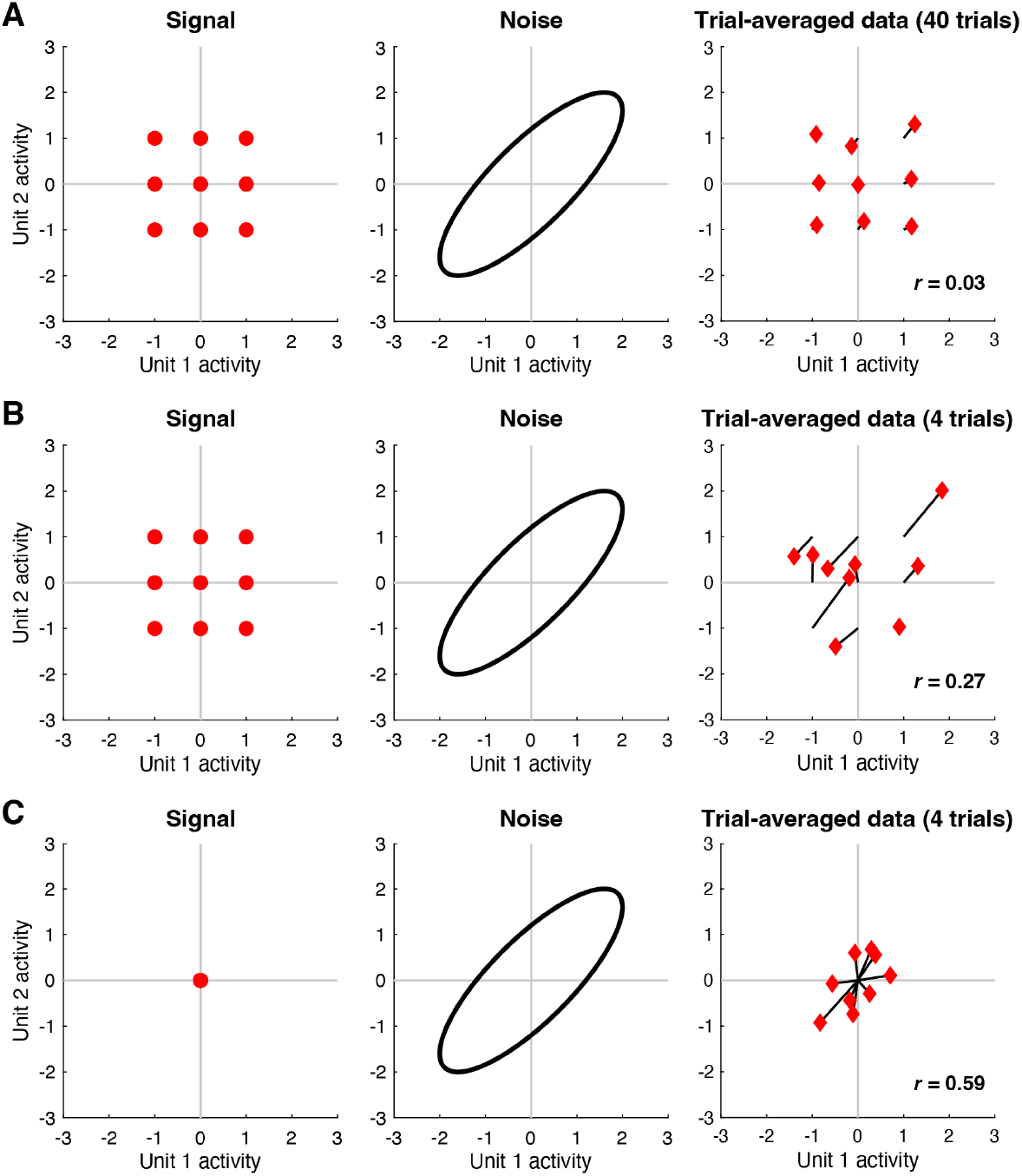
Trial averaging is insufficient for removing the effects of noise. Here we perform simulations to illustrate how noise correlations persist after trial averaging (code available at https://osf.io/fc589). *A*, In this simulation, responses to 9 conditions are measured from 2 units. The left shows the signal, i.e. responses in the absence of noise. The middle shows the noise, i.e. trial-to-trial response variability for a fixed condition; the noise is drawn from a zero-mean multivariate Gaussian distribution (ellipse indicates a Mahalanobis distance of 2). The right shows responses averaged across 40 trials for each condition (black lines join the trial average to the corresponding signal). *B*, Same as panel A except that 4 trials per condition are used. *C*, Same as panel B except that the signals associated with the 9 conditions are all set to zero.

In this paper, we propose an analysis technique for disentangling signal and noise covariance in neural response measurements. Our approach, termed generative modeling of signal and noise (GSN), builds and fits a model of the signal and noise components of measured multivariate neural responses. The model is *generative* in the sense that the process by which measurements are generated is explicitly modeled, and the model is *distributional* in the sense that it attempts to characterize how neural responses are distributed across conditions. (This latter characteristic contrasts with tuning-based models that attempt to characterize how neural responses vary as a function of specific properties of experimental conditions.) First, we lay out the principles underlying GSN and validate GSN through a series of simulations with a known ground truth. In conducting these simulations, we also compare the performance of GSN to that of several related techniques. Next, we demonstrate the application of GSN to visually evoked functional magnetic resonance imaging (fMRI) responses in the publicly available Natural Scenes Dataset (Allen et al., 2022). This provides intuition for how GSN fares on empirical brain data and highlight ways in which GSN can be leveraged within computational neuroscience. Finally, we use the example data to illustrate how GSN can be used to improve the results of principal components analysis. Specifically, by disentangling signal and noise, GSN provides a cleaner estimate of the signal in the data and its properties (eigenspectra and dimensionality).

While elements of the statistical components comprising GSN can be found in prior work (Duan et al., 2023; Ledoit and Wolf, 2004; Pospisil and Bair, 2021a; Pospisil and Pillow, 2024; Schäfer and Strimmer, 2005; Stringer et al., 2019; van Bergen and Jehee, 2021; Yatsenko et al., 2015), novel contributions of the present work include integrating principles and techniques into a clearly articulated framework, developing an algorithm and associated code toolbox for optimally fitting the GSN model, and demonstrating several ways in which GSN may be useful for neuroscience applications. The code used in this paper is available at https://osf.io/wkyxn/, and the code toolbox implementing GSN is available at https://github.com/cvnlab/GSN/.

## Results

### Generative signal and noise modeling approach

Consider the general situation where responses are measured from a set of units (e.g., voxels, neurons, channels) to several experimental conditions (e.g. stimuli) and several trials are collected for each condition. The core idea of the generative signal and noise (GSN) approach is to model each response as reflecting the sum of a sample drawn from a multivariate distribution associated with signal (defined as the response to different conditions in the absence of noise) and a sample drawn from a multivariate distribution associated with noise (defined as trial-to-trial response variability for a fixed condition). We assume the noise distribution is zero-mean and assume the noise sample is independent of the signal sample. We allow the signal and noise distributions to have potentially different covariances.

A schematic illustrating GSN is shown in **Figure 2**. This schematic depicts a situation in which responses are measured from two units to 40 conditions with three trials per condition. Panel A shows the ground-truth signal distribution. Red dots are samples from the distribution and indicate noiseless responses to the 40 conditions. One of the dots is highlighted in blue, marking an example condition. Panel B shows the ground-truth noise distribution. Blue x’s indicate three samples from the distribution; these are noise samples associated with the example condition. Panel C shows the data distribution, whose mean and covariance are equal to the sum of the means of the signal and noise distributions and the sum of the covariances of the signal and noise distributions, respectively. The red x’s indicate the measured responses (obtained as the sum of signal and noise), with the blue x’s highlighting the responses associated with the example condition. Overall, panels A–C illustrate how signal and noise distributions give rise to observed measurements.

**Figure 2.**
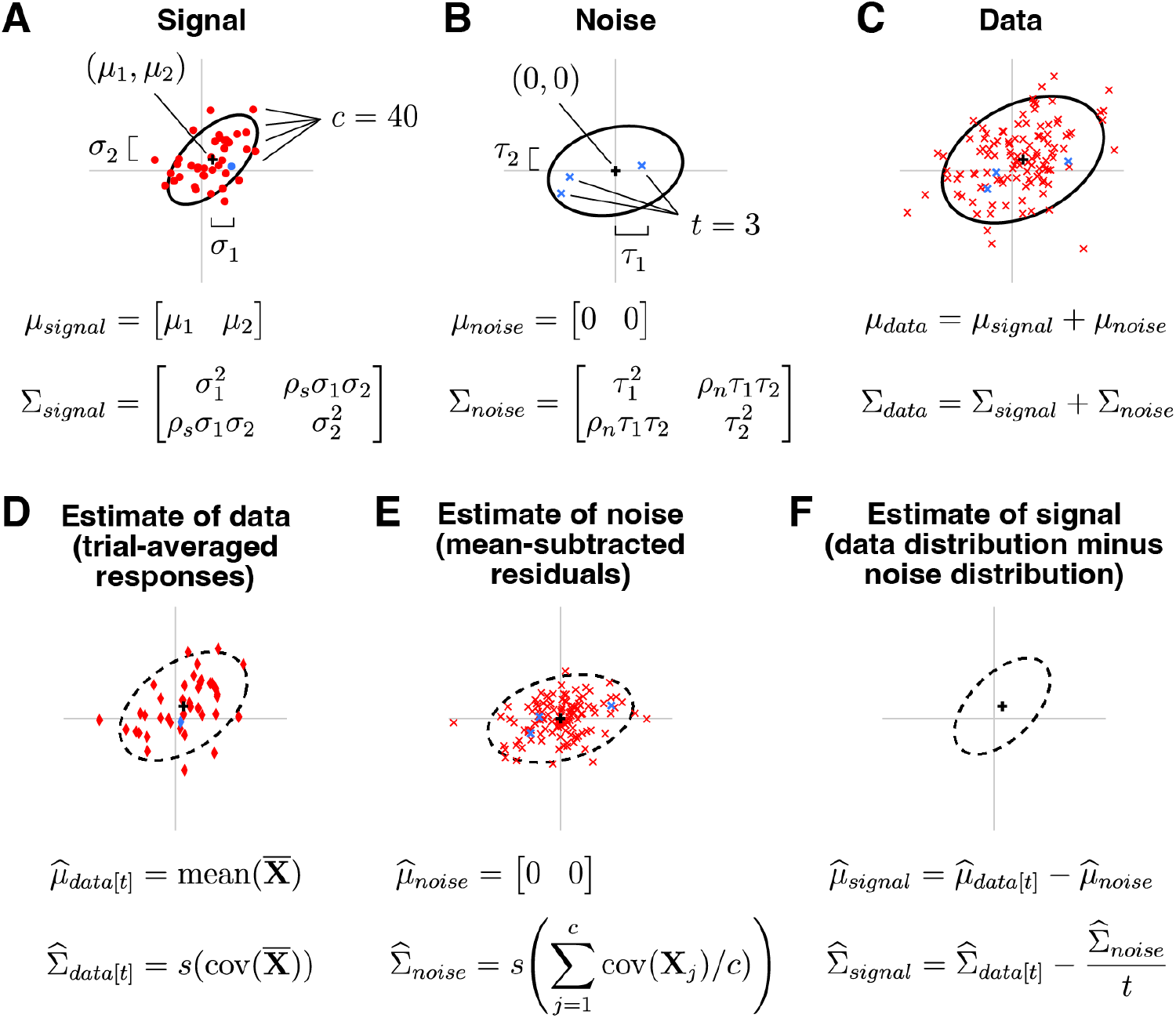
Schematic of GSN. Here we depict an example involving *n* = 2 units, *c* = 40 conditions, and *t* = 3 trials per condition (code available at https://osf.io/7k2m5). In each plot, the black cross and black ellipse indicate the mean and spread (Mahalanobis distance of 2) of a multivariate Gaussian distribution. For definitions of symbols, please see *Methods. A*, Signal. The signal indicates responses to different conditions in the absence of noise and is modeled as a multivariate distribution. *B*, Noise. The noise indicates trial-to-trial variability for a given condition and is modeled as a zero-mean multivariate distribution. *C*, Data. The data are modeled as the sum of a sample from the signal distribution and a sample from the noise distribution. *D*, Estimate of data distribution. Given a set of measured responses, we compute trial-averaged responses and estimate the mean and covariance of these responses, yielding the estimate of the data distribution. *E*, Estimate of noise distribution. We compute the covariance of responses to each condition and average across conditions, yielding the estimate of the noise distribution. *F*, Estimate of signal distribution. We subtract the estimated parameters of the noise distribution from the estimated parameters of the data distribution, yielding the estimate of the signal distribution.

The core challenge in GSN is estimating the unknown signal and noise distributions given a set of measurements. The basic procedure that we propose is illustrated in panels D–F. Responses are averaged across trials and the mean and covariance of the trial-averaged responses are computed, as shown in panel D (red diamonds indicate trial-averaged responses; the blue diamond corresponds to the example condition). This procedure yields the estimate of the data distribution. After subtracting the mean response from the original non-trial-averaged responses to each condition, the covariance of the residuals is computed and then averaged across conditions, as shown in panel E (red x’s indicate the residuals; blue x’s indicate the residuals associated with the example condition). This yields the estimate of the noise distribution. Finally, the parameters associated with the noise distribution are subtracted from the parameters associated with the data distribution, as shown in panel F. This is the key step that corrects for the noise that persists after trial averaging (see **Figure 1**), and yields the estimate of the signal distribution. In order to ensure positive semi-definite covariance estimates, the full procedure is more complicated than what is presented here (please see *Methods* for details).

### Validation of GSN through ground-truth simulations

GSN attempts to determine the signal and noise distributions that underlie a set of measured responses. To help validate GSN, we performed ground-truth simulations involving 10 units whose ground-truth signal and noise distributions have specific structure (**Figure 3**). For the signal distribution, each unit was set to have a variance of 1, and units 1 through 5 were given positive correlation (*r* = 0.5; covariance = 0.5). For the noise distribution, each unit was set to have a variance of 2, and units 4 through 8 were given positive correlation (*r* = 0.5; covariance = 1).

**Figure 3.**
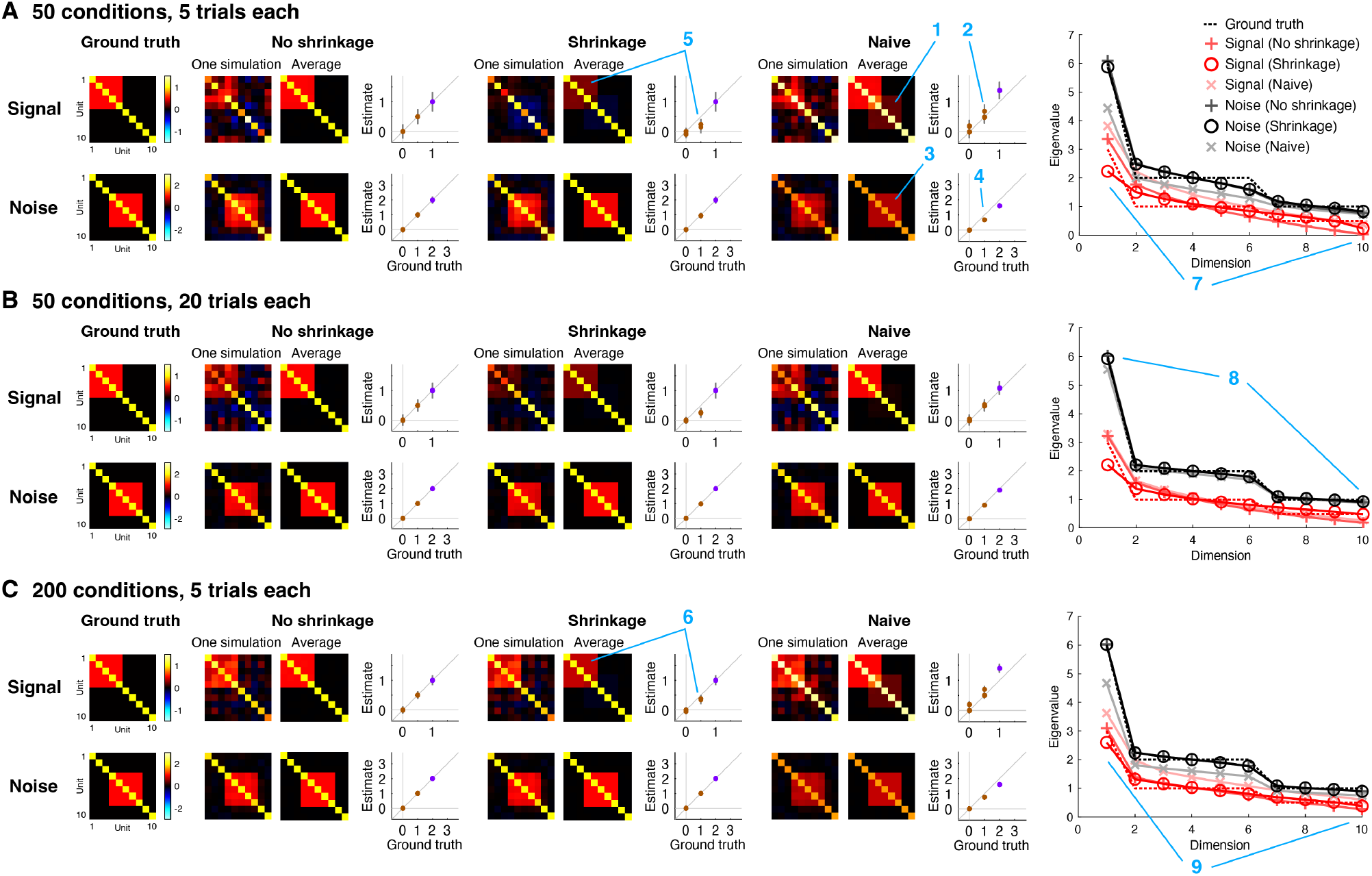
Estimation of signal and noise distributions. Here we show results of simulations that assess how well GSN estimates the signal and noise distributions that underlie a set of measurements (code available at https://osf.io/5uskr). All simulations involve 10 units whose responses are generated as the sum of a sample from a signal distribution and a sample from a noise distribution. Both distributions are multivariate Gaussian with zero mean but have different covariances (as depicted). For different combinations of number of conditions (samples from the signal distribution) and number of trials (samples from the noise distribution for each condition), we perform 1,000 simulations. In each simulation, we generate responses and analyze the resulting data using three different methods: ‘Naive’ refers to simple heuristic methods for estimating signal and noise covariance (see main text), ‘No shrinkage’ is the GSN method with standard covariance estimation, and ‘Shrinkage’ is the GSN method with shrinkage-based covariance estimation. Blue number labels highlight specific aspects of the results that are discussed in the main text. *A–C*, Detailed inspection of results for specific condition and trial numbers. In the scatter plots, purple and brown dots indicate diagonal and off-diagonal elements of the covariance matrix, respectively, and error bars indicate standard deviation across simulations. At the far right are plots of the eigenspectra (mean across simulations) produced by the three methods, as well as the ground-truth eigenspectra.

To gain insight, we plot detailed inspections of the performance of different methods for recovering ground-truth signal and noise distributions (**Figure 3**). First, consider the performance of naive methods for signal and noise estimation (‘Naive’). For signal estimation, the naive method is to simply average responses across trials and compute the sample covariance of the trial-averaged data (we refer to this method as ‘Signal (Naive)’). We see that this method incurs upward bias in the estimated signal covariance values; this can be observed in the qualitative image plots as the seeping of the noise covariance into the signal estimate (panel A, location 1) and in the quantitative scatter plots as dots lying above the line of unity (panel A, location 2). The bias is due to the fact that although trial averaging reduces noise, the trial-averaged data are still influenced by noise (Pospisil and Pillow, 2024). Thus, it is critical for an estimation procedure to account for this persistent noise. For noise estimation, the naive method is to simply remove the mean response for each condition, aggregate the residuals across conditions, and then proceed to covariance estimation (we refer to this method as ‘Noise (Naive)’). We see that the naive method for noise estimation incurs downward bias in the estimated noise covariance values; this can be observed in the image plots (panel A, location 3) and the scatter plots (panel A, location 4). The reason for this bias is that the naive method fails to account for the reduced degrees of freedom in the de-meaned responses: aggregating de-meaned responses across conditions involves using *ct*− 1 in the denominator of the calculation of covariance, whereas the correct approach is to use *t*− 1 in the denominator of the calculation of covariance for each condition, which is equivalent to a final denominator (after pooling) of *c*(*t*−1)=*ct*−*c*. Thus, the denominator is inflated in the naive method, leading to downward bias in the estimated covariance values.

We now proceed to the GSN method for signal and noise estimation. One version of GSN is coupled with standard covariance estimation (‘No shrinkage’), providing estimates of signal covariance (referred to as ‘Signal (No shrinkage)’ and of noise covariance (referred to as ‘Noise (No shrinkage)’). These estimates are unbiased (dots in the scatter plots lie on the line of unity) but suffer from high variance (error bars indicating standard deviation across simulations are large). A second version of GSN is coupled with shrinkage-based covariance estimation (‘Shrinkage’), providing estimates of signal covariance (referred to as ‘Signal (Shrinkage)’) and of noise covariance (referred to as ‘Noise (shrinkage)’). These estimates have reduced variance (brown dots indicating off-diagonal elements have smaller error bars), but are biased (the brown dots lie below the line of unity). Notice that the amount of bias is larger in scenarios with low amounts of data (e.g., panel A, location 5) than in scenarios with high amounts of data (e.g., panel C, location 6).

Besides assessing how well the different methods estimate covariance, we can also assess how well the different methods estimate eigenspectra. We observe that the sample covariance tends to underestimate dimensionality. This is most visible in the estimation of signal covariance when the number of conditions is small (panel A, location 7, red +’s). By incorporating shrinkage (panel A, location 7, red circles), the match to the ground-truth eigenspectrum is improved (panel A, location 7, red dashed line). Notice that the difference between the two methods diminishes in situations where a relatively large number of samples is available, such as estimation of noise covariance (panel B, location 8) or when the number of conditions is increased (panel C, location 9). Finally, consistent with earlier observations, we see that naive signal estimation produces eigenvalues that are too high (pink x’s; reflecting the seeping of the noise covariance into the signal covariance estimate) and that naive noise estimation produces eigenvalues that are too low (gray x’s; reflecting the lack of compensation for the reduced degrees of freedom).

We summarize the overall performance of the different methods by plotting ground-truth recovery of covariance as a function of number of trials (**Figure 4A**) and number of conditions (**Figure 4B**). In plotting these results, we also show the performance of an alternative method that is often used to estimate signal covariance (Pospisil and Pillow, 2024; Stringer et al., 2019). This method involves computing covariance of responses across independent splits of a given set of data (we refer to this method as ‘Split-half’). The intuition underlying the Split-half method is that the signal is expected to repeat across splits, whereas the noise is not expected to do so. Finally, we show results for additional scenarios beyond the simple idealized scenario depicted in **Figure 3**. In these additional scenarios (**Figure 4C**), we use randomly generated signal and noise covariances and explore the impact of varying the number of units and varying the dimensionality of the signal and noise distributions (see *Methods* for details).

**Figure 4.**
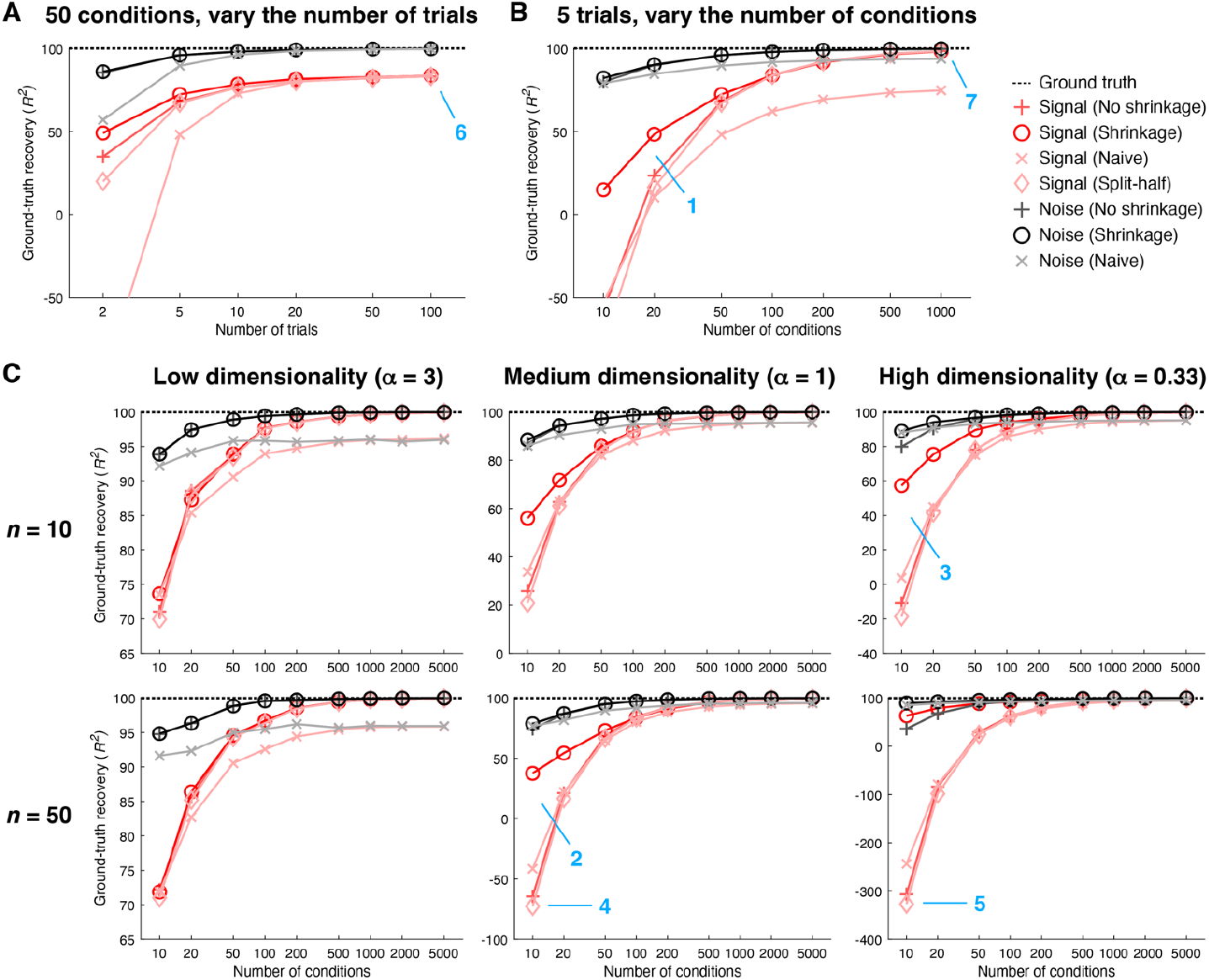
Ground-truth recovery of covariance. Here we quantify how well different methods recover signal and noise covariance (code available at https://osf.io/5uskr and https://osf.io/3yvtg). Performance is quantified using coefficient of determination (*R*^2^) with respect to values in the upper triangle of the covariance matrix (including the diagonal). The ‘Split-half’ method involves computing covariance across independent splits (trials) of the data. *A–B*, Recovery performance for the simple scenario illustrated in **Figure 3**. We vary the number of trials while holding the number of conditions fixed at 50 (panel A), and we vary the number of conditions while holding the number of trials fixed at 5 (panel B). Markers indicate the mean across 1,000 simulations. *C*, Recovery performance for a set of scenarios in which the number of units is varied (rows) and the dimensionality of the signal and noise is varied (columns). In these scenarios, signal and noise eigenspectra are governed by the power-law function *d*^−*α*^ where *d* is the 1-indexed dimension number and *α* is an exponent parameter. We fix the number of trials at 5 and vary the number of conditions. Markers indicate the mean across 50 simulations.

We find that in general, the noise distribution is easier to estimate than the signal distribution. This makes sense since all samples contribute to estimating the noise distribution, whereas only the mean of the samples associated with a condition contribute to estimating the signal distribution. We also see that across the board, the shrinkage method performs better than or as well as the other methods, with larger improvements in low-data regimes (e.g. panel B, location 1). This is consistent with the idea that although the No shrinkage method converges to the correct covariance when results are averaged across a large (infinite) number of simulations (i.e. it is unbiased), in individual simulations the Shrinkage method produces more accurate results than the No shrinkage method. Moreover, the benefit of shrinkage is especially pronounced in scenarios with high dimensionality (e.g., panel C, locations 2 and 3). This reflects the fact that non-regularized covariance estimates tend to underestimate dimensionality and shrinkage enables covariance estimates to become less correlated and hence higher-dimensional.

Notably, the Split-half method performs very similarly to the No shrinkage method. This makes sense from a theoretical standpoint: noise is expected to average out when computing covariance across splits and neither method incorporates shrinkage. However, notice that the Split-half method does systematically slightly underperform the No shrinkage method at low numbers of conditions (e.g., panel C, locations 4 and 5). One reason this may be the case is that the Split-half method is sensitive to stochastic sampling issues: results are dependent on exactly which trials are placed into each split, and performance presumably suffers unless one averages over all possible splits (which may be computationally impractical).

A final observation is that the limiting factor for accurate estimation appears to be the number of conditions available. In the simple scenario (as illustrated in **Figure 3**), if we fix the number of conditions at 50, even if we greatly increase the number of trials, ground-truth recovery of the signal reaches a plateau that is lower than 100% (**Figure 4**, panel A, location 6). This reflects the fact that although additional trials are helpful for reducing noise in the responses to individual conditions, the quality of signal covariance estimation is still limited by the number of samples drawn from the signal distribution. In contrast, if we fix the number of trials to 5, as we increase the number of conditions, ground-truth recovery of the signal approaches 100% (panel B, location 7). In other words, even if the number of trials per condition is low, we can achieve accurate recovery of signal and noise distributions as long as we sample a sufficient number of conditions. This means that when designing an experiment in which we can either sample more trials per condition or sample more conditions, if one’s goal is to accurately estimate signal and noise covariance, it is more important to sample many conditions than to sample many trials per condition. Alternatively, if one’s goal is to obtain accurate estimates of the mean response to each condition, sampling more trials per condition is more important.

### Recovery of effective dimensionality and power-law exponent

There has been increasing interest in studying the dimensionality of neural representations (e.g. (Canatar et al., 2023; Jazayeri and Ostojic, 2021; Pospisil and Pillow, 2024; Stringer et al., 2019)). A simple and useful metric of dimensionality is effective dimensionality (ED) (Del Giudice, 2021), which summarizes the distribution of eigenvalues in an eigenspectrum with a single number. A different metric stems from modeling eigenspectra using a power-law function (Stringer et al., 2019). Power-law functions are straight lines in log-log space; hence, a convenient metric of dimensionality is the slope of a line corresponding to a power-law function in log-log space, which is equivalent to the exponent of the power-law function. An interesting open question is how well the signal and noise estimates provided by GSN enable these dimensionality metrics to be recovered. We therefore augmented our simulations with additional analyses. Whereas our earlier analyses (in **Figure 4**) quantify how well a given method recovers signal and noise covariance values in terms of variance explained (*R*^2^), here we sought to quantify how well a given method recovers ground-truth values for ED and the power-law exponent. Hence, there are two differences in the evaluations: one difference concerns the quantity being recovered (covariance values vs. summary metrics of covariance structure), and the other difference concerns the evaluation criterion (variance explained vs. absolute difference between the ground-truth value and the estimated value).

For the same scenarios shown in **Figure 4C**, we calculated the ED and power-law exponent associated with the ground-truth signal and noise covariances, and compared these ground-truth values to the estimates provided by different methods (**Figure 5**). For each data point (reflecting a particular combination of scenario, number of conditions, and number of trials), we performed 50 simulations and computed the average estimate obtained across simulations. This allows us to investigate whether we can expect a given method to recover, on average, the ground-truth ED and power-law exponent, or whether the method exhibits bias. In conducting these analyses, we also included for comparison the performance of two methods that have been recently proposed for estimation of signal eigenspectra: cvPCA (Stringer et al., 2019), which is based on computing variance across independent splits of a given set of data, and MEME (Pospisil and Pillow, 2024), which is based on optimizing an eigenspectrum model to match the moments of the signal eigenspectrum estimated from a given set of data (see *Methods* for details).

**Figure 5.**
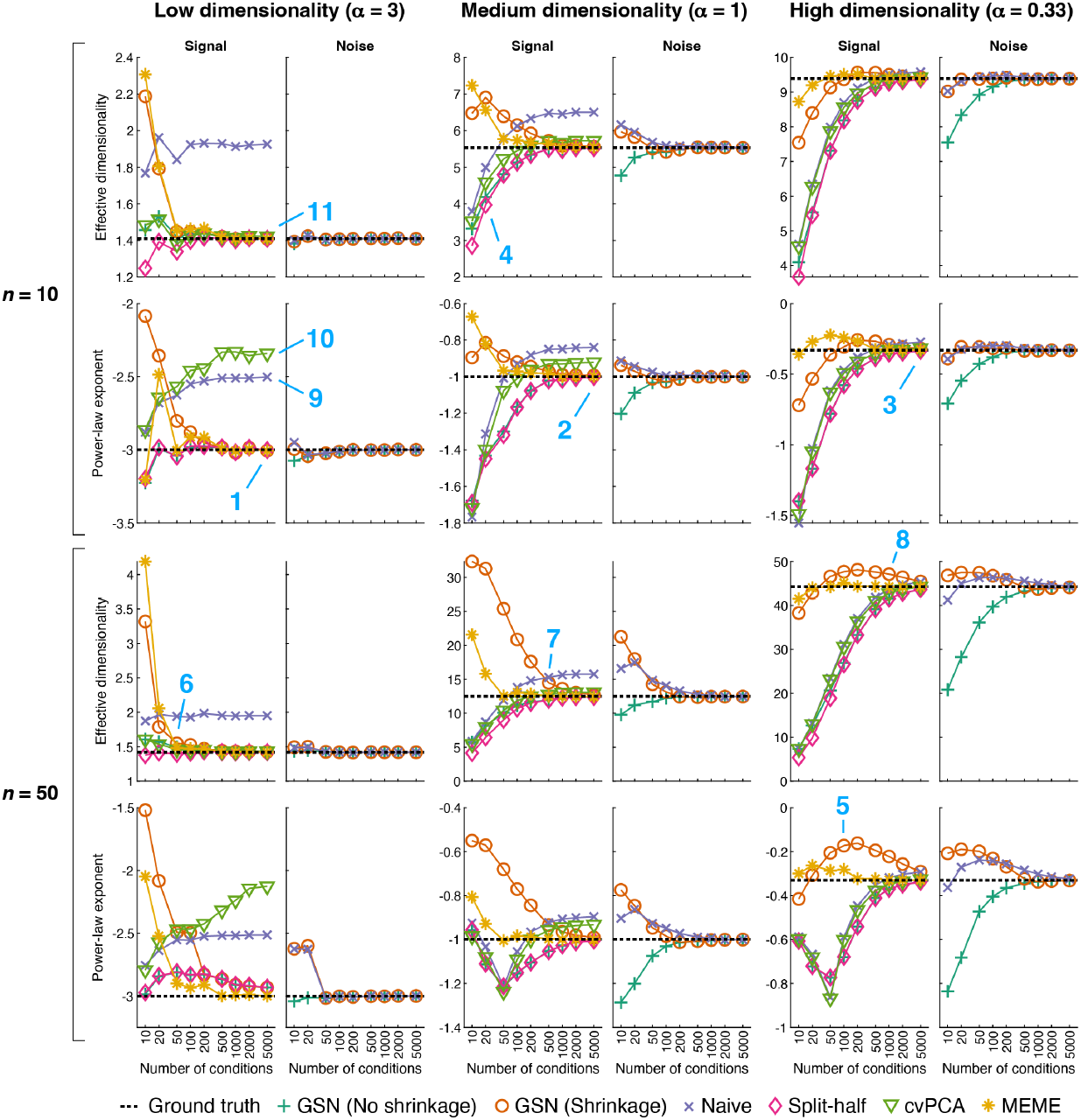
Ground-truth recovery of effective dimensionality and power-law exponent. Here we quantify how well different methods recover two summary metrics of signal and noise eigenspectra: effective dimensionality and power-law exponent (code available at https://osf.io/3yvtg). Recovery performance is plotted for the same scenarios shown in **Figure 4C**. The cvPCA method estimates the signal eigenspectrum by projecting two splits of a given set of data onto principal components (PCs) and calculating the dot product between the two sets of projections obtained for each PC. The MEME method estimates the signal eigenspectrum by estimating signal eigenmoments from a given set of data and then adjusting the parameters of an eigenspectrum model to match the estimated eigenmoments. Markers indicate the mean across 50 simulations, and the horizontal dotted line indicates the ground-truth value. Note that the Split-half, cvPCA, and MEME methods do not provide estimates for the noise (and are therefore not plotted).

The simulation results show a variety of interesting observations. On the whole, several of the methods perform reasonably well: GSN (No shrinkage), GSN (Shrinkage), Split-half, and MEME all provide estimates that converge towards ground-truth values at large number of conditions (e.g. locations 1, 2, 3). Hence, these methods provide the means to track and recover dimensionality of different scenarios. However, all methods exhibit bias at low numbers of conditions (e.g. location 4), and biases sometimes persist even for numbers of conditions that may seem relatively large, such as 100 (e.g. location 5). This underscores the point that collecting sufficient amounts of data is critical for accurately estimating dimensionality. As might be expected, the necessary amount of data scales with the dimensionality of the scenario being characterized (e.g. compare locations 6, 7, 8). Also, similar to earlier observations (see **Figure 4**), it is easier to recover properties of noise than properties of signal.

While ranking methods is tricky given the high complexity of the patterns of results, we venture some general conclusions. The worst performing methods are Naive and cvPCA, as they tend to exhibit high bias even for large numbers of conditions (e.g. locations 9, 10). The Split-half and GSN (No shrinkage) methods perform at about the same level (echoing results in **Figure 4**), and converge towards ground-truth values at modest rates. In contrast, GSN (Shrinkage) converges towards ground-truth values more rapidly. The best performing method is MEME which exhibits the fastest convergence to ground-truth values. However, the MEME method comes with limitations, including assumptions about the shape of the eigenspectrum (see Discussion).

It is interesting to note that qualitatively different patterns of results can be found for ED and power-law exponent. For example, performance of a given method can be relatively poor for power-law exponent (location 10) but relatively good for ED (location 11). We interpret this as simply reflecting the fact that different metrics emphasize different aspects of eigenspectra. Another observation is that the various methods perform well even when the number of units increases 5-fold from 10 to 50. Presumably, what is relevant is not only the raw number of units but also the dimensionalities of the signal and noise that are distributed across the units (which might be low).

To further explore the generality of our conclusions, we performed simulations for an additional scenario involving biologically realistic signal and noise covariances. Results are generally similar, except for complications related to the recovery of power-law exponent (see **S1 Figure** for details). Finally, we caution that while our simulations indicate reasonable performance across a range of settings, our simulations are not comprehensive (e.g., we did not specifically vary the relative magnitudes of signal and noise) and practitioners may wish to perform simulations matched to the system being studied if precise values are critical.

### Application of GSN to empirical data

#### Signal and noise covariance estimates

We demonstrate the application of GSN to empirical data taken from the 7T fMRI Natural Scenes Dataset (NSD) (Allen et al., 2022). NSD consists of human brain responses to over 70,000 visually presented natural scenes distributed across eight participants. Each image is presented up to three times to a given participant. This limited number of presentations reflects the prioritization of sampling a large number of distinct images over sampling a large number of trials per image (see also (Stringer et al., 2019)). As such, NSD can be viewed as an especially challenging dataset for methods that seek to accurately disentangle signal from noise.

As an illustrative example, we extracted responses from right hemisphere fusiform face area subdivision 1 (FFA-1) in one participant (Participant 1), yielding 330 vertices × 10,000 images × 3 trials. As a pre-processing step, we normalized the responses associated with each vertex to have zero mean and unit variance. We then performed GSN on these data, yielding estimates of signal and noise covariance (**Figure 6**).

**Figure 6.**
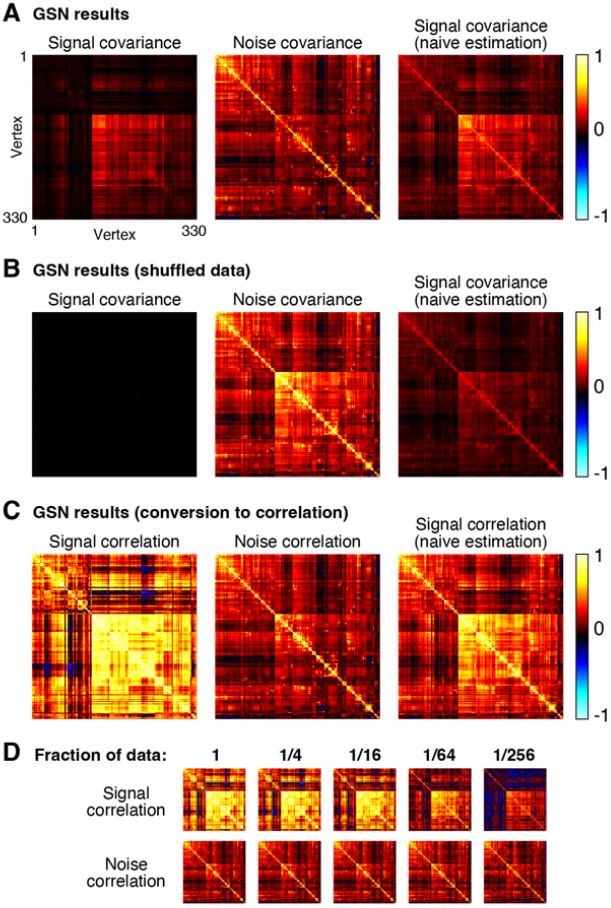
Application of GSN to example fMRI data. Here we demonstrate the application of GSN to example data from FFA-1 (330 vertices × 10,000 images × 3 trials) (code available at https://osf.io/yxrsp). *A*, Signal and noise covariance estimates. In addition to GSN outputs (first and second columns), we show results from naive estimation of signal covariance which involves simply calculating the covariance of trial-averaged data (third column). *B*, Results for shuffled data. As a control, we shuffled responses across all images and trials and re-analyzed the data. *C*, Conversion to correlation units. The results of panel A are re-plotted after converting covariance to correlation units. *D*, Estimates as a function of amount of data. We varied the fraction of images to which GSN is applied (e.g. 1/16 corresponds to 625 of 10,000 images being used). This was done such that data subsets were mutually exclusive of one another.

A number of observations can be made from the results. First, notice that the magnitude of the noise is generally larger than the magnitude of the signal (panel A, compare diagonal of noise covariance with diagonal of signal covariance). The fact that response measurements contain large trial-to-trial variability even when holding the experimental manipulation (stimulus) constant is typical in fMRI and many other measurement techniques. Second, we observe that the covariance structure of the noise is different from that of the signal, though there are some similarities (panel A, compare middle image with left image). A naive method that averages responses across trials yields covariance structure (panel A, right image) that is a mixture of signal covariance and noise covariance, since trial averaging reduces but does not eliminate noise. Third, as a control, if we fully shuffle responses across images and trials, we see that values in GSN’s estimated signal covariance become very low (panel B, left image). This makes sense since after shuffling, we do not expect to find reliable differences in responses across images. In contrast, the naive method fails to produce a good signal covariance estimate: even though there are no reliable differences in responses across images, trial averaging does not fully suppress the noise and the noise covariance seeps into the signal covariance estimate (panel B, right image).

For visual comparison, we show covariance estimates after conversion to correlation units (panel C). One motivation for this conversion is to ensure that each unit contributes equally to subsequent analyses of the covariance estimates. Prominent differences between covariance and correlation are observed, reflecting the fact that there are substantial variations in signal-to-noise ratio across vertices (vertices with low signal strength are only weakly visible in the covariance matrices and become more visible in the correlation matrices). Finally, by applying GSN to different subsets of the data (panel D), we see that signal and noise can be reliably estimated in this dataset. For example, compare the signal and noise correlation estimates obtained using 1/4th of the data to those obtained using 1/16th of the data (these reflect two mutually exclusive subsets of the data). Reliable estimation is especially notable given that the dataset involved only three trials for each stimulus. Of course, in the limit of very low amounts of data (panel D, rightmost columns), estimation quality starts to suffer and we start to see strong influence of the shrinkage bias pulling off-diagonal elements towards zero.

Although GSN does not require nor assume Gaussian distributions, if the signal and noise distributions are indeed Gaussian, then the mean and covariance parameters estimated by GSN are sufficient for a full characterization of a given dataset. Curious about the nature of the distributions in NSD, we performed inspections of the example data shown in **Figure 6**. In these inspections, we compare histograms of the empirical data to histograms of synthesized data that are generated using parameters of the GSN model coupled with the assumption of Gaussian signal and noise distributions (**S2 Figure**). We find a high level of similarity for a histogram of trial-averaged responses (which helps focus on signal) and a histogram of mean-subtracted residuals (which helps focus on noise), suggesting that the signal and noise indeed have Gaussian-like distributions.

#### Eigenspectra of signal and noise

Principal components analysis (PCA) is a widely used method for dimensionality reduction and data visualization (Greenacre et al., 2022). Using the empirical data, we conducted several analyses that demonstrate the benefits of GSN for PCA. The first analysis (**Figure 7A**) pertains to eigenspectra, which are important as they indicate the amount of variance explained by different principal components. For each of the eight NSD participants, we computed the eigenspectrum of the covariance of the data after trial averaging (cyan lines); this represents a naive analysis in which responses are averaged across trials to reduce noise. We also computed the eigenspectrum of the signal covariance (red lines) and noise covariance (black lines) as estimated by GSN. To make the results directly comparable to the results of the naive analysis, we scaled the noise covariance by 1/3 (since trial averaging is expected to reduce the variance of the noise by a factor equal to the number of trials). Finally, we calculated the effective dimensionality associated with each of the three eigenspectra (numbers above each plot).

**Figure 7.**
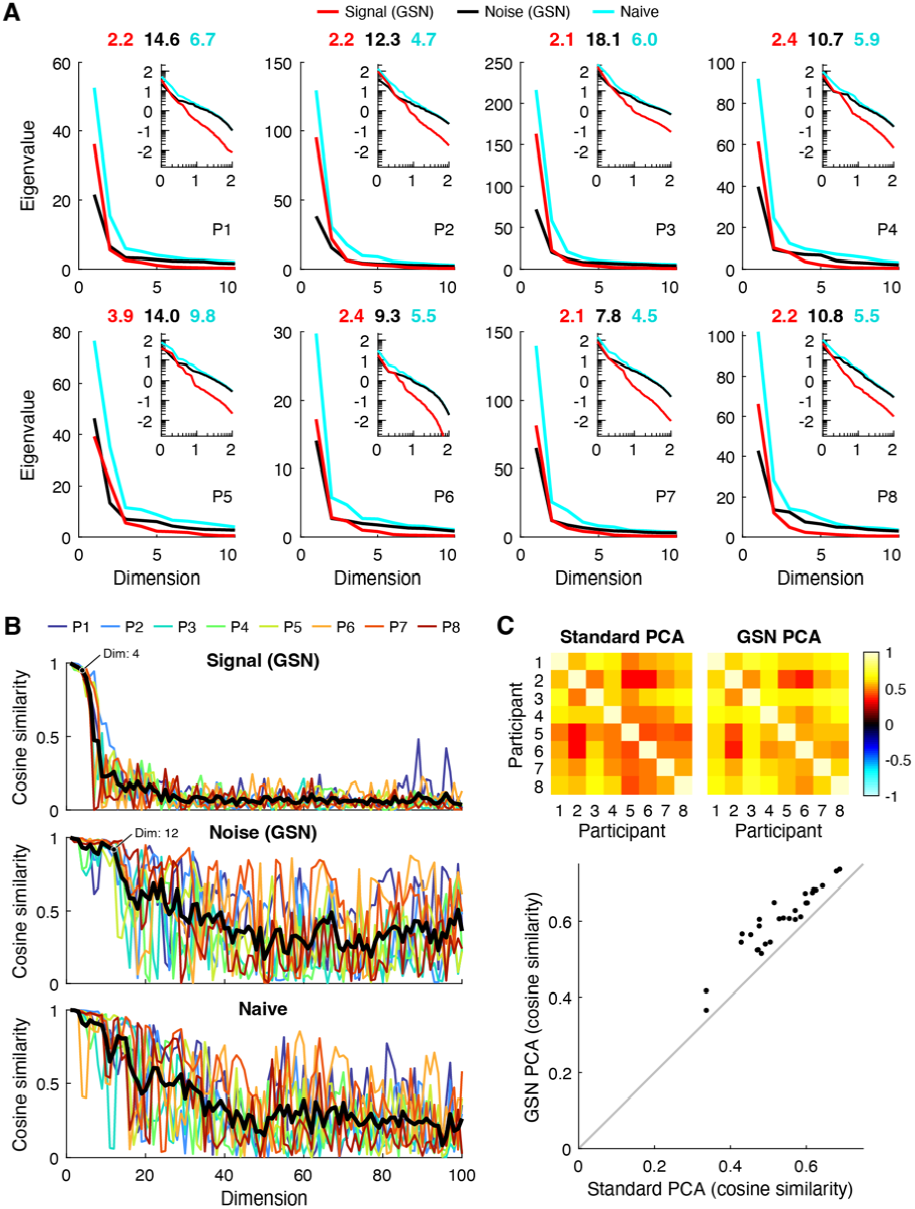
GSN disentangles signal and noise in principal components analysis (PCA). Here we use PCA to analyze the results of GSN as applied to FFA-1 (code available at https://osf.io/f34bc). *A*, Eigenspectra. For each of the eight participants (P1–P8), we plot the eigenspectra of the signal and noise as estimated by GSN (‘Signal (GSN)’, ‘Noise (GSN)’), as well as the eigenspectrum of the trial-averaged data (‘Naive’). The main plots show results on a linear scale for up to the first 10 dimensions; the insets show results on a base-10 log-log scale for up to the first 100 dimensions. Numbers above each main plot indicate the effective dimensionality of the three eigenspectra. *B*, Split-half reliability of principal components. The cosine similarity between corresponding principal components from two split-halves of the data from each participant is plotted for up to the first 100 dimensions. The thick black line indicates the mean across participants. *C*, Across-participant consistency. A common set of 515 images were viewed three times each by all participants. For each participant, we computed the projections of trial-averaged responses to these 515 images onto either (i) the first principal component of the covariance of the trial-averaged data (‘Standard PCA’) or (ii) the first principal component of the signal covariance estimated by GSN (‘GSN PCA’). The cosine similarity of these projections between each pair of participants is shown.

We find that the three eigenspectra exhibit distinct patterns. In terms of overall magnitudes, eigenvalues are slightly higher for the signal than they are for the noise and are highest for the naive analysis. This is consistent with the interpretation that after averaging across 3 trials, the total variance contributed by signal is slightly higher than the total variance contributed by noise, and that the trial-averaged data have high total variance due to contributions from both signal and noise. In terms of how quickly eigenvalues fall off (independent of their magnitudes), we see that the eigenspectrum of the signal falls off relatively quickly and has low effective dimensionality (between 2–4). This indicates that the coding of natural scenes in FFA-1 is low-dimensional (at least as measured in NSD). In contrast, we find that the eigenspectrum of the noise falls off more slowly and has higher effective dimensionality than the signal (between 9–18). This is most evident in the inset log-log plots, which show more clearly what occurs at high numbers of dimensions. Finally, we find that the eigenspectrum of the trial-averaged data falls somewhere in the middle, with a moderate effective dimensionality (between 4–10). Overall, these results illustrate how GSN separates signal and noise components in a set of data and enables the researcher to study their separate properties. The separation of noise from signal is important, as it compensates for the fact that in empirical data, noise corrupts the dimensionality of the measured signal (Del Giudice, 2021).

#### Reliability of principal components

A second analysis pertains to the reliability of the principal components derived from the data. We randomly split the images from each participant into halves, performed PCA separately on the two split-halves, and then computed the cosine similarity of principal components across the split-halves. Results are shown both for the signal and noise as estimated by GSN as well as for the naive trial-averaged data (**Figure 7B**). We find that the principal components of the signal are highly reliable across split-halves for approximately the first 4 dimensions, and that the principal components of the noise are highly reliable for approximately the first 12 dimensions (see labeled points). Beyond these numbers of dimensions, reliability levels are substantially lower, which makes sense given that the amount of variance associated with the higher dimensions is very small (see **Figure 7A**). The principal components of the trial-averaged data also exhibit reasonably high levels of reliability. However, the reliability levels decrease gradually, making it difficult to decide the number of highly reliable dimensions.

One peculiar observation is that reliability values for the noise and the trial-averaged data fluctuate, but on average stay elevated, over a large range of dimensions (20–100). We suggest that this could be due to the fact that the eigenvalues in these higher dimensions are roughly equal in magnitude, making the ordering of the principal components somewhat arbitrary and subject to estimation error. In such a scenario, corresponding principal components across split-halves are not likely to match but might incidentally match on occasion. Finally, notice that the reliability pattern for the trial-averaged data looks approximately like a mixture of the reliability patterns for the signal and the noise. This is consistent with the interpretation that the data is a mixture of signal and noise and that GSN successfully decomposes the data into these constituent components.

#### Denoising of PCA results

The third and final analysis seeks to validate the signal and noise identification provided by GSN. In short, how do we know that GSN is successfully estimating and removing the influence of noise? Here, we can leverage the notion that signal, not noise, is expected to generalize across participants (Charest et al., 2018). We reasoned that if GSN successfully separates signal from noise in each participant, then signal properties—specifically, the coding of natural scenes—should exhibit improved consistency across participants compared to the trial-averaged data. This is because the trial-averaged data is expected to contain the residual effects of noise, and many types of noise are expected to be idiosyncratic to each participant (e.g., the effects of head motion on fMRI responses is likely unrelated to the coding of natural scenes). But how can we compare participants? Given the variability of the size and shape of FFA-1 across participants (the number of vertices is not even the same), comparing principal components across participants is not straightforward. However, we can compute the projections of responses to natural scenes onto principal components, and these projections should be comparable across participants insofar that there is some degree of commonality in the representation of natural scenes across participants.

In accordance with our approach for assessing across-participant consistency, we computed trial-averaged responses for a common set of 515 images that were viewed by all participants, and then projected these responses onto the top principal component of the signal covariance estimated by GSN. For comparison, we also projected the responses onto the top principal component of the trial-averaged data. The results show that the projections for GSN are substantially more consistent across participants than the standard analysis (**Figure 7C**). This implies that GSN is successfully reducing the influence of noise on principal components derived from the data, and that the principal components derived by GSN better reflect the underlying coding dimensions in the brain that are shared across humans. As a sanity check, we visually inspected the stimulus images that drive variance along the direction of the top principal component (**S3 Figure**); this reveals that the presence of faces appears to be the dominant factor, consistent with prior studies (Grill-Spector et al., 2017; Kanwisher et al., 1997).

Finally, we show results of our PCA analyses for additional brain regions V1, hV4, and PPA (**S4 Figure**). Our main observations replicate, including lower dimensionality for the signal compared to the noise, high within-participant reliability of the first several signal PCs and noise PCs, and higher across-participant consistency of trial-averaged response projections onto PC1 for GSN PCA than for standard PCA. In addition, we find that the dimensionality of the signal is substantially higher in V1 (mean across subjects: 5.6) and hV4 (mean across subjects: 4.6) than it is in FFA-1 (mean across subjects: 2.4), whereas the signal dimensionality is comparable in PPA (mean across subjects: 2.5). These variations in dimensionality across brain regions are a desirable outcome, as they are consistent with the idea that GSN is able to track and recover different dimensionality levels. More generally, these results indicate that GSN can aid the investigation of representational differences across the brain.

## Discussion

In this paper, we have described a simple generative model that characterizes the contributions of signal and noise to a set of neural response measurements. We developed a method for fitting this model, implemented this method in a code toolbox, and demonstrated the method on ground-truth simulations and empirical data. We showed four main results. First, we showed that naive approaches to estimating signal covariance (i.e. trial averaging) and estimating noise covariance (i.e. aggregating residuals) are inaccurate (**Figures 3–6**). A key insight is that simply computing trial-averaged responses is insufficient to eliminate noise: the result will invariably contain a mixture of both signal and noise covariance. Second, we confirmed that the GSN method works as expected, with ground-truth recovery performance improving with larger numbers of trials and conditions (**Figure 4**). Third, we performed simulations directly comparing GSN to alternative methods for signal estimation (including split-half analyses, cvPCA, and MEME), and found that GSN is competitive with these methods (**Figure 5**). Fourth, we showed how GSN can be exploited to improve principal components analysis (PCA). Specifically, GSN decomposes a set of data into signal and noise distributions, each of which has its own eigenspectrum and eigenvectors. These distributions can be analyzed separately, for example, with respect to dimensionality (**Figure 7A**) and reliability (**Figure 7B**). Furthermore, isolating the signal distribution leads to principal components that have improved generalizability across participants (**Figure 7C**).

### Novel contributions of the present work

Elements of GSN can be found in prior work, including using repeated trials to separate signal and noise in neural responses (Henriksson et al., 2015; Pospisil and Bair, 2021a; Pospisil and Pillow, 2024; Stringer et al., 2019) and the use of shrinkage for covariance estimation (Ledoit and Wolf, 2004; Schäfer and Strimmer, 2005; van Bergen and Jehee, 2021; Yatsenko et al., 2015). We note, in particular, that the formulation of the GSN model is fairly close to the approach described in a recent pre-print in the statistics literature (Duan et al., 2023). Overall, the work presented here is best viewed as an applied statistics paper, one that selects and consolidates statistical ideas and designs methods for application to a specific scientific domain (neural response measurements). The primary novel contributions of the present work are the integration of techniques into a clearly articulated framework, developing an algorithm for optimally fitting the GSN model under the constraint of positive semi-definite covariance estimates, and demonstrating specific examples of how GSN could be useful in neuroscience applications. In addition, we provide a code toolbox for easy application of GSN.

### Relationship to other approaches

From a statistical perspective, GSN bears some similarity to probabilistic principal components analysis (PPCA) (Ghojogh et al., 2021; Roweis, 1997; Tipping and Bishop, 1999). PPCA is a special case of factor analysis, and models the data as the sum of the combination of latent factors and a noise term. However, a key difference between PPCA and GSN is that PPCA assumes that the noise is isotropic (i.e., the noise has the same variance and is uncorrelated across units), whereas GSN does not make this assumption. Instead, GSN exploits the fact that neural response measurements usually involve multiple trials per condition, and estimates the noise structure instead of assuming it to be isotropic. Another difference is that PPCA typically comes with the presumption that the latent variables have lower dimensionality than the original data, whereas GSN does not necessarily involve dimensionality reduction.

Signal and noise correlations have been studied in the computational neuroscience literature using a variety of approaches. Here, we discuss a few approaches closely related to GSN. The approach used in (Triplett et al., 2020) involves building a model of calcium imaging data that simultaneously characterizes both evoked activity (signal) and spontaneous activity (noise). The model is generative in nature, similar to GSN. A difference is that the approach involves a number of modeling choices that are specific to the signal and noise characteristics present in calcium imaging data. Incorporating modality-specific details may enhance statistical efficiency and interpretability. In contrast, GSN has a different philosophical goal of providing a general-purpose framework for signal and noise estimation that rests on minimal assumptions. Another generative modeling approach, TAFKAP, was introduced by (van Bergen and Jehee, 2021, 2018) in the context of developing improved decoding methods for fMRI data. This approach, like GSN, estimates both signal covariance and noise covariance. However, the modeling of signal proceeds quite differently in TAFKAP than GSN. In TAFKAP, the response of each unit to the experimental conditions is fit using a specific tuning curve model—for example, in (van Bergen and Jehee, 2021), a weighted sum of basis functions is used to model the orientation tuning of each unit. GSN takes a different approach: instead of attempting to estimate the signal (noiseless response) to each condition, GSN attempts to estimate only the distribution of the signal across conditions. An advantage of the GSN approach is that it avoids the need to specify (and thus does not depend on) a tuning curve model, thereby providing more generality. Moreover, if a tuning curve model is used, there is a risk that model failures (either due to model misspecification or imperfections in model fitting) may corrupt estimates of the noise (assuming noise is estimated from model residuals) (Wilson and Gardner, 2023). However, a disadvantage of the GSN approach is that it requires condition repeats to estimate the noise, whereas in TAFKAP, noise can be estimated based on residuals of the model fit. Another disadvantage is that the lack of an explicit tuning model in GSN implies that further analysis steps must be carried out in order to incorporate GSN into decoding analyses.

### Comparison to cvPCA and MEME

A recent paper (Stringer et al., 2019) proposed a method termed ‘cross-validated PCA’ (cvPCA) that seeks to quantify signal (stimulus-related variance) in neural response measurements, similar to GSN. The method involves splitting a dataset into halves (where the halves contain different trials for the same set of conditions), performing PCA on one half, projecting the responses in each half onto the estimated PCs, and then computing covariance across the projections from each half as an estimate of signal variance. The underlying logic is that noise is not expected to covary across halves, whereas the signal is expected to do so. Similar to GSN, the cvPCA method leverages repeated trials to infer what is related to the experimental manipulations (signal) and relies on a model in which the total variance in a dataset is equal to the sum of signal variance and noise variance. However, the two methods differ substantially in the procedures by which estimates are obtained.

Recent work (Pospisil and Pillow, 2024) has pointed out that the PCs estimated in cvPCA are influenced by noise and are therefore not identical to the true underlying signal PCs. This fact degrades the accuracy of the signal components estimated by cvPCA, and leads to biased estimates of the signal eigenspectrum (Pospisil and Pillow, 2024). Motivated by these concerns, the authors propose the MEME (minimize eigenmoment error) method to deliver improved estimates of the signal eigenspectrum. Specifically, MEME first calculates unbiased estimates of the moments of the signal eigenspectrum, assumes a parametric model for the signal eigenspectrum, and then optimizes parameters of the model to minimize the error between the eigenspectrum moments achieved by the model and the estimated moments of the signal eigenspectrum.

In this paper, we performed simulations that directly compare the performance of GSN, cvPCA, and MEME with respect to recovery of effective dimensionality and power-law exponent (see **Figure 5** and **S1 Figure**). We observed substantial bias in cvPCA results, consistent with recent reports (Pospisil and Pillow, 2024). GSN performs better, converging towards ground-truth values with increasing amounts of data. MEME performs the best, with even faster convergence. However, a major limitation of MEME is that it assumes a parametric form for the signal eigenspectrum. In our main set of simulations (**Figure 5**), for simplicity we considered only scenarios where the ground-truth signal eigenspectrum fully conformed to the form assumed by our MEME implementation (specifically, a single unbroken power-law function). Deviations from the assumed form are expected to lead to degraded performance from MEME. Indeed, it is possible that deviation from the assumed form is responsible for the poor performance of MEME in recovering power-law exponent in the biologically realistic scenario (**S1 Figure**). While allowing break points in the power-law function may help ease the constraints of MEME, doing so increases complexity and may lead to instability in parameter optimization. As the user must hand-pick initial guesses for parameter values, it might be challenging to come up with robust choices for initial parameter values for break points.

Overall, the cvPCA and MEME methods are similar in spirit to GSN in the sense of using repeated trials to separate signal and noise in neural response measurements. However, the former two methods are primarily focused on estimation of signal eigenspectra, whereas GSN takes a broader view in which the goal is to estimate full covariance matrices (including both the eigenspectrum and eigenvectors) for the signal and the noise. As such, GSN supports a wider array of subsequent analyses of signal and noise properties. It might be possible to use eigenspectrum estimates from cvPCA or MEME to produce improved estimates of full covariance matrices, but the originally described methods do not do so, which is why we do not compare to such hybrid or extended methods here.

### Other applications of GSN

Besides improving PCA and dimensionality estimation (as illustrated in **Figure 7**), GSN may aid in other applications not specifically covered in this paper. One important application is the estimation of noise ceilings for computational models (Lage-Castellanos et al., 2019; Pospisil and Bair, 2021b). Since noise imposes limits on the maximum amount of variance that can in theory be predicted on the basis of experimental events (e.g. sensory stimuli), obtaining accurate estimates of the noise ceiling is critical for assessing model performance. GSN provides explicit models of the distributions of signal and noise, and can be used to estimate noise ceilings for the responses of individual units (see Methods in (Allen et al., 2022)) as well as noise ceilings for multivariate measures, such as representational dissimilarity matrices (see Methods in (Conwell et al., 2022)). Having principled methods to compute univariate and multivariate noise ceilings is critical in efforts to compare deep neural network models of brain data at scale (Cichy et al., 2019; Conwell et al., 2022; Schrimpf et al., 2020; Willeke et al., 2022).

Another application relates to research programs where noise itself is of intrinsic interest, often hypothesized to perform functions relevant to neural computation (e.g., (Bays, 2014; Dinstein et al., 2015; Ecker et al., 2014; Keeley et al., 2020; Ma et al., 2006; Orbán et al., 2016; Stein et al., 2005; van Bergen et al., 2015)). The GSN approach facilitates the study of noise by decomposing datasets into signal and noise, providing researchers with two distinct entities that can be separately measured, characterized, manipulated, compared with one another, and related to brain function. Isolating the separate contributions of signal and noise to response measurements may help enrich our understanding of how response variability contributes to the function of neural systems and whether and how noise and signal interact.

### Limitations of GSN and future directions

GSN rests on the assumption that noise is additive and independent of the signal. The assumption of independence simplifies estimation and enables efficient use of data: even though the example dataset in this paper included only three trials per image, pooling estimates of noise covariance across images enabled robust noise covariance estimates (see **Figure 6D**). The extent to which the additive and independence assumptions accurately characterize fMRI responses is an important open question. For example, a recent study provided evidence that noise magnitude and noise correlations in fMRI data decrease during task states (Ito et al., 2020). However, it is clear that the additive and independence assumptions do not strictly hold for spiking data. Spike trains exhibit Poisson-like proportionality between the mean firing rate and the variance of firing rate across trials (Tolhurst et al., 1983), and this proportionality may depend upon stimulus statistics (Festa et al., 2021). Moreover, multiplicative-type noise has been observed in which firing rates in neural populations are collectively scaled (Goris et al., 2014; Lin et al., 2015; Liska et al., 2022). Finally, evidence that noise depends on the stimulus has been shown for neurons in the retina (Franke et al., 2016; Zylberberg et al., 2016). A direction for future work would be to relax the assumptions of GSN to accommodate a larger range of settings.

Another potential limitation of GSN is that it may require a large number of samples for accurate estimation of signal and noise distributions. We observed that a relatively large number of conditions is required to accurately estimate the signal covariance (see **Figure 4B**). In addition, although pooling of noise covariance estimates across conditions can achieve robust estimation of noise (see **Figure 6D**), if one wishes to explore the possibility that the noise distribution may depend on the experimental condition, large numbers of trials for each condition may be required. Future research might investigate practical data requirements for a diverse range of experimental scenarios. A third limitation is that GSN in its current form provides just a point estimate of model parameters. If one is interested in the reliability of parameter estimates, it may be possible to extend GSN using bootstrapping or Bayesian techniques to obtain confidence intervals or posteriors for model parameters.

There is a sizable statistical literature on techniques for covariance matrix estimation (reviewed in (Fan et al., 2016)). Our proposed method for estimating covariance only incorporates shrinkage to improve estimation accuracy. This is a mild prior and is expected to improve out-of-sample generalization compared to an unbiased estimator. Within the technique of shrinkage, there are variants that can be tried such as deriving the optimal level of shrinkage analytically or using different shrinkage targets (Ledoit and Wolf, 2022, 2004; Schäfer and Strimmer, 2005). If one is willing to make stronger assumptions, there are other approaches that could achieve more efficient covariance estimates. Such approaches include banding and tapering (Bickel and Levina, 2008a), thresholding (Bickel and Levina, 2008b), and methods that impose low-rank structure (Pourahmadi, 2013; Yatsenko et al., 2015). In addition, one could seek to model covariance in terms of one or more structured covariance components (Pourahmadi, 2013; Triplett et al., 2020; van Bergen and Jehee, 2021; Yatsenko et al., 2015). This type of approach can improve estimation efficiency, but its utility depends on the accuracy of the assumed covariance components. If one is willing to make an explicit distributional assumption, one can apply Bayesian inference (e.g. (Leonard and Hsu, 1992)), which allows regularization through the prior. Finally, one could apply robust statistics (den Haan and Levin, 1997) to improve estimation. These various methods for covariance estimation could be easily incorporated into the GSN framework by simply replacing the shrinkage estimators that we use.

Finally, an important direction for future research is to devise methods for distinguishing different sources of noise. Neural noise (true variability in neural activity) is fundamentally distinct from instrumental noise (e.g. electrical noise), physiological noise (e.g. noise related to respiration and the cardiac cycle), and motion-related noise (e.g. motion of the head). Without specific modeling of these various noise sources, it remains unknown how much of the noise observed in a set of measurements is due to neural noise. Developing methods to identify non-neural noise and isolate neural noise will presumably lead to improved insights into the nature of noise and how it may support brain function.

## Methods

### The GSN method

#### Basic framework

GSN is a multivariate generalization of the univariate framework that we previously proposed for modeling signal and noise in responses of individual units (Allen et al., 2022). Consider the general situation in which responses are measured from a set of *n* units (e.g., voxels, neurons, channels) to *c* conditions (e.g., different stimuli) and this process is repeated for *t* trials per condition (we assume *t* > 1). In this scenario, response measurements have a dimensionality of *n* units × *c* conditions × *t* trials. The scenario is multivariate in the sense that there exist multiple units and we are attempting to model the joint distribution across all units. Our broad goal is to formally characterize the distribution of signal, i.e., the average expected response to each given condition, and the distribution of noise, i.e., trial-to-trial variability in the response to each given condition.

For the purposes of modeling, we assume that the signal and the noise are independent and additive and that each is characterized by some underlying multivariate distribution. We propose the following model:

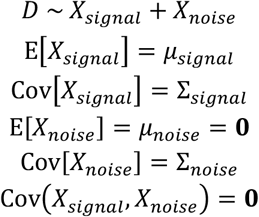

where *D* is an *n*-dimensional random variable indicating the responses of the *n* units on each trial (1 × *n*), *X*_*signal*_ is the signal component of the data with mean *µ*_*signal*_ (1 × *n*) and covariance Σ_*signal*_ (*n × n*), *X*_*noise*_ is the noise component of the data with mean *µ*_*noise*_ (1 × *n*) and covariance Σ_*noise*_ (*n × n*), and **0** indicates a matrix of zeros. In other words, the response on each trial is modeled as the sum of a random sample drawn from a signal distribution (which represents the noiseless response to some condition) and a random sample drawn from a noise distribution (which represents the noise that accompanies the response). The noise is assumed to be zero-mean. See **Figure 2A–C** for a visual illustration.

The modeling approach we describe is generative in the sense that we are characterizing the process by which measurements are generated (specifically, the data for each trial are modeled as a random draw from the multivariate distribution associated with *D*). We therefore refer to the approach as *generative modeling of signal and noise (GSN)*. Note that a complete generative model requires choosing specific forms for the distribution of signal and the distribution of noise; a simple choice is the multivariate Gaussian distribution (see **S2 Figure**).

### Algorithm for estimating model parameters

The core challenge in GSN is estimating the parameters of the signal and noise distributions. We propose a method based on the observation that the sum of two independent random variables has a mean that is equal to the sum of the means of the distributions associated with the variables and a covariance that is equal to the sum of the covariances of the two distributions. Hence, we can write:

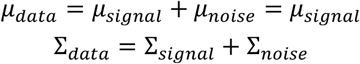

where *µ*_*data*_ and Σ_*data*_ indicate, respectively, the mean (1 × *n*) and the covariance (*n × n*) of the measurement variable *D*. For simplicity, we have used notation that acts as if each trial involves a fresh draw from the signal distribution. However, in typical practice, several trials are measured for each condition and the draw from the signal distribution is the same for each of these trials. To account for this, we average responses across the available *t* trials before estimating the data distribution. Since trials are independent, averaging is expected to reduce the covariance of the noise by a factor of *t*. Hence, we can write the following for the distribution of the trial-averaged data:

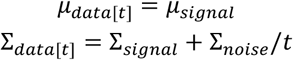

where *µ*_*data*[*t*]_ indicates the mean of the multivariate distribution that describes trial-averaged data (1 × *n*) and Σ_*data*[*t*]_ indicates the covariance of this distribution (*n × n*).

The general approach of GSN is to estimate the mean and covariance of the noise, estimate the mean and covariance of the trial-averaged data, and then subtract the noise covariance estimate (scaled by 1/*t*) from the trial-averaged data covariance estimate to obtain an estimate of the signal (see schematic in **Figure 2D–F**). However, because it is possible that the obtained estimate of signal covariance may be not positive semi-definite (especially in scenarios with limited data or low signal-to-noise ratio), a more sophisticated approach is necessary. To meet this challenge, we develop a mathematical formalism in which we use a weighted sum-of-squares approach to find positive semi-definite matrices for signal and noise covariance estimates that are as close as possible to the estimates derived directly from the data (details in **S5 Appendix**). This turns out to be a convex optimization problem that can be solved using an iterative approach, as we detail below.

The following is a step-by-step algorithm for GSN (*performgsn*.*{m,py}*):

1. Start with a set of neural response measurements ***X*** (*n* units × *c* conditions × *t* trials). Let ***X***_*j*_ denote the responses measured for condition *j*, arranged as a 2D matrix (*t* trials × *n* units). Let 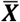 denote trial-averaged responses, arranged as a 2D matrix (*c* conditions × *n* units).
2. To estimate the noise distribution, calculate the covariance of responses separately for each condition, average the covariances across conditions, and then shrink the result. This yields an initial estimate of the noise covariance, which we refer to as 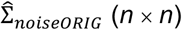:

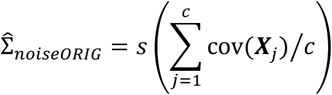

where 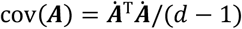 computes sample covariance using Bessel’s correction, 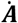 indicates ***A*** with its columns centered around zero, *d* is the number of rows in ***A***, and *s*() is a shrinkage procedure (see *Shrinkage-based regularization of covariance* below). Intuitively, we are quantifying unit-to-unit covariation around the mean response to each condition, pooling covariance estimates across conditions to improve accuracy, and then using shrinkage to further improve accuracy. Since we might update our estimate of the noise covariance later in the algorithm (if the signal covariance estimate turns out to be not positive semi-definite), we use 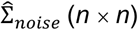 to refer to our current estimate of the noise covariance:

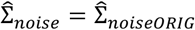

We assume that the noise distribution is zero-mean (i.e., the expected value of the noise for each unit is zero):

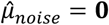

where 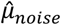 is the estimated noise mean (1 × *n*).
3. To estimate the data distribution (i.e., the distribution that characterizes the measured data), take the trial-averaged responses and then estimate mean and covariance, again applying shrinkage to improve accuracy of covariance estimation:

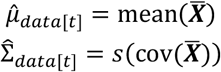

where mean() indicates column-wise mean, 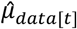 is the estimated data mean for the case of averaging across *t* trials (1 × *n*), and 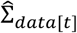 is the estimated data covariance for the case of averaging across *t* trials (*n × n*). Notice that 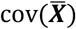 is the naive estimate of signal covariance that is obtained after simply trial averaging.
4. To estimate the signal distribution, subtract the current estimate of the noise distribution scaled by 1/*t* from the estimated data distribution:

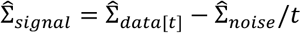

where 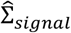 is the estimated signal covariance (*n × n*). Additionally:

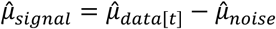

where 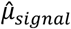 is the estimated signal mean (1 × *n*).
5. If the signal covariance estimate is positive semi-definite, we are done. Otherwise, proceed to Step 6.
6. Repeat until convergence:
  6.1 Calculate an updated estimate of the signal covariance:

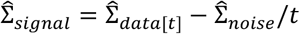

Ensure the signal covariance estimate is positive semi-definite by finding the nearest positive semi-definite matrix:

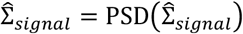

where PSD() is a method for finding the nearest symmetric positive semi-definite matrix to a given square matrix (details below).
  6.2 Calculate an updated estimate of the noise covariance:

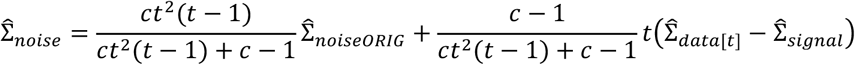

This calculates a weighted average of two possible estimates of the noise covariance: the first is the estimate based on the covariance of the mean-subtracted residuals (as calculated in Step 2), while the second is the estimate based on the subtraction of the signal distribution from the data distribution. The weights reflect the number of samples that inform each of the two estimates (see **S5 Appendix** for details). Ensure the noise covariance estimate is positive semi-definite by finding the nearest positive semi-definite matrix:

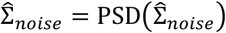
  6.3 If the correlation between the current and previous signal covariance estimates and the correlation between the current and previous noise covariance estimates are both greater than 0.999, stop. Otherwise, return to Step 6.1.

Convergence of the algorithm is guaranteed because the optimization problem is biconvex. In practice, convergence typically takes just a few iterations. For instance, in the execution of the set of simulations underlying **Figure 5**, the maximum number of iterations required by GSN was 3 (corresponding to the case where two updates are calculated beyond the initial estimates).

### Theoretical analysis of the GSN estimates

The proposed algorithm for GSN can be viewed as providing least-squares estimates of signal and noise covariance under the constraint that the estimates are positive semi-definite. Note that as long as the true signal and noise covariances are positive semi-definite, our estimators for them are consistent. This is because the original estimates 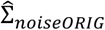 and 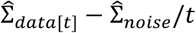 are consistent estimators of Σ_*noise*_ and Σ_*signal*_, respectively. For large enough datasets, the probability that the initial signal covariance estimate leaves the positive semi-definite cone thus goes to 0. Hence, our iterative estimates converge to the same values as the original estimates. For the same reason, our iterative estimates inherit all asymptotic properties of the original estimates. In particular, they are asymptotically unbiased and efficient, and the rate of convergence is the same as for the original estimates.

The primary advantage of our iterative estimates is that they are guaranteed to produce positive semi-definite (and, thus, valid) covariance matrices. Having valid covariance matrices is critical, as it is required for performing many subsequent analyses (such as PCA). While the constraint of positive semi-definiteness can be viewed as introducing bias, the true covariance matrices are known to be positive semi-definite. Hence, requiring positive semi-definiteness is warranted. Furthermore, due to the convexity of the cone encompassing positive semi-definite matrices, the projection of estimates onto this cone always reduces the distances (errors) to the true covariance matrices (see **S5 Appendix**).

### Shrinkage-based regularization of covariance

An appealing feature of computing sample covariance using Bessel’s correction is that the covariance values are unbiased estimates of the true covariance values. However, when the number of observations is small relative to the number of variables (in our case, when the number of trials or conditions is small relative to the number of units), the sample covariance is unstable and hence inaccurate. Moreover, the sample covariance may have an eigenspectrum that suffers from bias. To improve accuracy of covariance estimation, the GSN algorithm incorporates shrinkage (in Steps 2 and 3), a well-established method for regularizing covariance estimates (Chen et al., 2010; Daniels and Kass, 2001; Ledoit and Wolf, 2004; Schäfer and Strimmer, 2005). Specifically, the off-diagonal elements of the sample covariance are scaled towards zero, reflecting the prior that variables are generally expected to be uncorrelated. The goal of shrinkage is to introduce some amount of bias in order to reduce estimation variance and achieve a covariance estimate that is closer to the ground-truth covariance. Note that shrinking towards a diagonal matrix tends to increase the rank (dimensionality) of the covariance estimate. Also, note that shrinkage is not a requirement of the GSN approach and can be omitted if desired (using the flag <wantshrinkage>).

To perform shrinkage, we calculate:

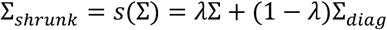

where Σ is the sample covariance (*n × n*), λ is a shrinkage fraction in the range [0,1], Σ_*diag*_ is Σ with off-diagonal elements set to zero (*n × n*), and Σ_*shrunk*_ is the shrinkage-based covariance estimate (*n × n*). When the shrinkage fraction is 1, the sample covariance is preserved and no shrinkage is applied; when the shrinkage fraction is 0, full shrinkage is applied. Notice that in our formulation, the target towards which estimates are shrunk (Σ_*diag*_) contains the original sample variance estimates on the diagonal. This choice of target is referred to as Target D “diagonal, unequal variance” in (Schäfer and Strimmer, 2005). The reason for this choice of target is to avoid imposing bias on the variances associated with the variables.

To determine the amount of shrinkage to apply, we use a cross-validation approach (similar to that used in (van Bergen and Jehee, 2021; Yatsenko et al., 2015)) in which held-out data are used to evaluate likelihoods corresponding to covariance estimates at different levels of shrinkage. We opt for this computational approach, as opposed to analytical methods for setting the shrinkage level (Ledoit and Wolf, 2004; Schäfer and Strimmer, 2005), for increased transparency and to avoid reliance on assumptions. In our implementation (*calcshrunkencovariance*.*{m,py}*), we randomly split the available data into an 80% training set and a 20% testing set. In the case of noise estimation (Step 2), the data are split with respect to trials; in the case of data estimation (Step 3), the data are split with respect to conditions. The sample covariance of the training set is then calculated, different shrinkage fractions ranging from 0 to 1 in increments of 0.02 are applied, the average negative log likelihood of observations in the testing set is calculated for each shrinkage fraction, and the shrinkage fraction yielding the minimum negative log likelihood is selected. In this way, the procedure derives a balance between bias and variance (the procedure will impose just enough bias to mitigate the damaging effects of variance). Note that in the case of estimating the noise distribution, the mean response to each condition in the testing set is subtracted before evaluating likelihoods (in order to remove the signal).

Our implementation includes flexible options that allow the user to control the training/testing split (<leaveout>) as well as the specific shrinkage fractions evaluated (<shrinklevels>). In addition, the implementation includes an optional flag (<wantfull>) that enables a final step in which the selected shrinkage fraction is applied to the sample covariance of the full dataset (combining both the training and testing sets). This option improves estimation quality (since more data are used) at the expense of imposing slightly more shrinkage than is optimal (in theory, if more training data are available, then less shrinkage should be necessary).

We conducted simulations to confirm the validity of our shrinkage-based method for covariance estimation (**S6 Appendix**). These simulations also confirm that shrinkage reduces the bias present in the eigenspectrum of the sample covariance.

### Method for finding the nearest positive semi-definite matrix

To ensure valid covariance matrices, the GSN algorithm involves finding the nearest (in the sense of the Frobenius norm) symmetric positive semi-definite matrix to a given matrix (see PSD() in Steps 6.1 and 6.2). This is accomplished using the method proposed by Higham (Higham, 1988). Our implementation is as follows (*constructnearestpsdcovariance*.*{m,py}*):

1. Start with a given square matrix ***C***.
2. Ensure symmetry by updating ***C*** = (***C*** + ***C***^T^)/2.
3. Perform singular value decomposition to obtain ***C*** = ***USV***^T^.
4. Compute the approximating matrix 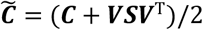.
5. If 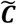 is not positive semi-definite (due to numerical precision issues), add a small multiple of the identity matrix (*ε****I***) to 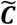 and restart the procedure starting from Step 3. We use *ε* = 10^−10^.

Note that this method is equivalent to performing an eigendecomposition of ***C*** and setting negative eigenvalues to zero.

### Additional analyses related to GSN

#### Conversion of covariance to correlation

When interpreting covariance matrices, it is often useful to convert the values to correlation units. Correlation is simply a version of covariance where the variances of each of the two variables have been normalized to one. We provide a function to convert covariance matrices to correlation units (*convertcovariancetocorrelation*.*{m,py}*). Our implementation divides each element of a given covariance matrix by the square root of its associated row-wise diagonal element and by the square root of its associated column-wise diagonal element. This conversion procedure is used in **Figure 6**.

#### Principal components analysis

The present study uses principal components analysis (PCA) as a means for interpreting the results of GSN. We perform PCA through eigendecomposition of a given covariance matrix:

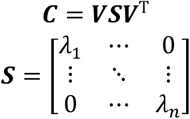

where ***C*** is a covariance matrix (*n × n*) associated with data in *n* dimensions, ***V*** is an orthonormal matrix (*n* × *n*) with unit-length eigenvectors in the columns, and ***S*** is a diagonal matrix (*n × n*) with eigenvalues along the diagonal in descending order (λ_1_ ≥ λ_2_ ≥ ⋯ ≥ λ_*n*_ ≥ 0). The eigenvectors are referred to as *principal components*; the sizes of the eigenvalues indicate the importance of the principal components; and the full set of eigenvalues is referred to as the *eigenspectrum*. A given data point (1 × *n*), expressed relative to the centroid of the data, can be projected onto the principal components, producing *scores* (1 × *n*). These scores are simply the coordinates of the data point in the rotated space defined by the principal components. Finally, a useful metric that summarizes the distribution of eigenvalues is effective dimensionality (ED) (Del Giudice, 2021):

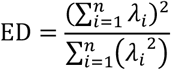

This metric ranges continuously from 1 to *n* and indicates the number of underlying dimensions in the data (specifically, the number of dimensions that results in an equivalent amount of entropy). Note that the metric shown above is just one of several possible metrics for ED (Del Giudice, 2021).

Depending on one’s goals, one might want to convert a covariance matrix to correlation units before computing the eigendecomposition. The motivation for this would be to ensure that all dimensions have equal influence (otherwise, dimensions with larger variances would tend to dominate the principal components). Indeed, in standard usage of PCA, it is generally recommended to *z*-score each dimension as a pre-processing step; this has the consequence that the covariance matrix will be in correlation units.

### Ground-truth simulations

We conducted ground-truth simulations to illustrate key concepts, test our code implementation, and evaluate the performance of different methods. All simulations involved generating synthetic response measurements based on multivariate Gaussian signal and noise distributions.

#### Data scenarios

We designed three types of data scenarios:

1. First, we created easy-to-interpret scenarios involving simple structure for signal and noise covariance. These scenarios involved 10 units, and are used in **Figures 3 and 4A** and **S6 Appendix**.
2. Second, we created a set of scenarios that systematically varied the number of units and the dimensionality of the signal and the noise. These scenarios are used in **Figures 4C and 5**. In these scenarios, the ground-truth signal and noise distributions were each zero-mean and had a covariance that was constructed by combining a randomly generated set of eigenvectors and a power-law eigenspectrum. Specifically, eigenspectra were governed by the power-law function 1/*d*^*α*^ = *d*^*α*^ where *d* indicates the 1-indexed dimension number and *α* indicates an exponent parameter. Scenarios involved either 10 or 50 units and either an exponent of *α* = 3 (low dimensionality), *α* = 1 (medium dimensionality), or *α* = 0.33 (high dimensionality) for the signal and noise, resulting in a total of six scenarios. For each number of units (10, 50), we generated random eigenvectors independently for the signal and the noise and held these eigenvectors constant across scenarios with different exponents.
3. Third, we created a biologically realistic scenario in which ground-truth signal and noise distributions were taken to be the empirical GSN signal and noise distribution estimates obtained for right hemisphere FFA-1 in Participant 1 as shown in **Figure 6**. This is, of course, somewhat provisional since it assumes that the estimates provided by GSN are reasonable. Nonetheless, the choice is justifiable since the goal of our simulations is to evaluate ground-truth recovery in simulated data (as opposed to making an inference about empirical data). The scenario involved 330 units, and is used in **S1 Figure**.

Each scenario was simulated using specific combinations of numbers of conditions (*c*) and numbers of trials (*t*). For each combination of *c* and *t*, multiple simulations were performed in order to average out incidental variability. The standard errors of results across simulations were sufficiently small and therefore are not shown in the figures.

#### Estimation methods

Given a set of response measurements generated in a simulation, we applied five different methods for estimating aspects of the signal and noise. The methods are as follows:

1. *GSN (No shrinkage)* - This is the GSN method coupled with standard covariance estimation.
2. *GSN (Shrinkage)* - This is the GSN method coupled with shrinkage-based covariance estimation.
3. *Naive* - For signal estimation, the naive method is to simply average responses across trials and compute the sample covariance of the trial-averaged data. For noise estimation, the naive method is to simply remove the mean response for each condition, aggregate the residuals across conditions, and then perform covariance estimation.
4. *Split-half* - This method refers to computing covariance across independent splits of a dataset as a means for signal covariance estimation, and has been previously used in the literature (Pospisil and Pillow, 2024; Stringer et al., 2019). Our implementation of the method is as follows. Given a set of response measurements (*n* units × *c* conditions × *t* trials), we randomly divide the trials into two equal splits (or nearly equal in the case of an odd number of trials), average responses across trials within each split, compute covariance across the splits, and then average the resulting covariance matrix with its transpose to ensure symmetry. Formally, the signal covariance estimate is given by 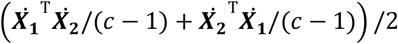 where ***X***_**1**_ and ***X***_**2**_ indicate trial-averaged responses arranged as a 2D matrix (*c* conditions × *n* units) for the two splits, respectively, and 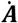 indicates ***A*** with its columns centered around zero. We perform 10 random splits of the trials, and average the signal covariance estimate across splits. We note that variants of the method are possible, including performing exhaustive trial splits (in the case of low numbers of trials) and calculating covariance across pairs of trials.
5. *cvPCA* - The cross-validated PCA (cvPCA) method is described in (Stringer et al., 2019), and delivers an estimate of the signal eigenspectrum. We start with the same preparation as described for the Split-half method: 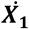 and 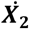 are centered, trial-averaged responses (*c* conditions × *n* units) for two splits of the data. We compute principal components (PCs) of the first split, project the responses in each split onto these PCs, and then compute the dot product between the two sets of projections obtained for each PC dimension. Formally, the signal eigenspectrum estimate is given by 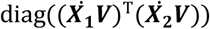 where ***V*** indicates the PCs (*n* units × *n* dimensions) obtained from the first split. We perform 10 random splits of the trials, and average the signal eigenspectrum estimate across splits. We note that other variants of the cvPCA method are possible, such as performing multiple iterations where responses to each condition are shuffled across the two splits (Stringer et al., 2019).
6. *MEME* - The minimize eigenmoment error (MEME) method is described in (Pospisil and Pillow, 2024), and delivers an estimate of the signal eigenspectrum. Given a set of response measurements (*n* units × *c* conditions × *t* trials), we randomly divide the trials into two equal splits (or nearly equal in the case of an odd number of trials) and average responses across trials within each split. We then apply the MEME method as implemented in the code provided at https://github.com/dp4846/meme_v1_bpl/blob/master/src/eig_mom.py (function *fit_broken_power_law_meme_W*). A high-level overview of the procedure is as follows. First, the user chooses a parametric model—specifically, a broken power-law function—for the eigenspectrum. Then, moments of the eigenvalues of the covariance matrix (i.e. eigenmoments) are estimated from the data. Finally, nonlinear optimization is used to optimize parameters of the model in order to minimize the squared error between the moments of the modeled eigenspectrum and the moments estimated from the data. We use the fitted model parameters returned by the code to reconstruct the estimate of the signal eigenspectrum. We perform 10 random splits of the trials, and average the signal eigenspectrum estimate across splits.

The MEME implementation requires specifying several hyperparameters: the number of eigenmoments to consider, a list of break points where the power-law function might be broken, and initial guesses for the power-law intercept and the slopes of the power-law segments. For our simulations, we make the following choices. First, we set the number of eigenmoments to consider to 5. Second, given that the ground-truth eigenspectra are exactly linear in log-log space in the main set of simulations (**Figure 5**), we do not use the MEME functionality for estimating breakpoints and instead use a single (unbroken) power-law function as the parametric model. Third, to give the MEME method the best possible chance for accurate estimation, we set the initial guesses for the slope and intercept of the power-law line to the ground-truth values. In our tests, we found that MEME results are generally robust to the choice of initial guesses (e.g., using generic values often gave good results); however, we noticed that results are more unstable when using initial guesses that are farther from the ground-truth values, suggesting that caution should be exercised when setting hyperparameters in real analysis contexts.

We evaluated the performance of the methods with respect to three different metrics. One metric is recovery of signal and noise covariance values. For a given method’s estimate of covariance (either of the signal or of the noise), the coefficient of determination (*R*^2^) between the estimated covariance values and the ground-truth covariance values was calculated, and the average *R*^2^ across simulations was computed. The second metric is recovery of effective dimensionality (ED). For a given method’s estimate of the eigenspectrum (either of the signal or of the noise), ED was computed, and the average ED across simulations was compared to the ED of the ground-truth eigenspectrum. The third metric is recovery of power-law exponent. For a given method’s estimate of the eigenspectrum (either of the signal or of the noise), a line was fit to the estimated eigenspectrum in log-log space (details below) and the slope of the line was recorded. The average slope across simulations was compared to the slope of a line fit to the ground-truth eigenspectrum.

Note that the Split-half method generates estimates of only the signal covariance, and is therefore evaluated only in terms of recovery of signal covariance, signal ED, and signal exponent. Also, note that the cvPCA and MEME methods generate estimates of only the signal eigenspectrum, and are therefore evaluated only in terms of recovery of signal ED and signal exponent.

### Line fitting method

To determine the power-law exponent corresponding to a given eigenspectrum, we fit a line to the eigenspectrum in log-log space where the *x*-axis corresponds to the 1-indexed dimension number and the *y*-axis corresponds to the eigenvalue. To ensure robust results across diverse simulations, we designed a heuristic procedure that appears to work well in practice. First, given an eigenspectrum of length *d*, we create a linear grid in log space from log(1) to log(*d*) using a granularity that is at least as fine as the separation between log(*d*–1) and log(*d*). This grid is transformed back to linear space and rounded to the nearest integer, producing a set of indices. The motivation for this rounding procedure (which is a method used in the code provided with (Stringer et al., 2019)) is to avoid interpolation of eigenvalues. Next, we define “good” eigenvalues as those that are greater than 0.001 of the maximum eigenvalue. This excludes very small, zero, and negative eigenvalues, all of which can degrade the quality of line fits. (If only one eigenvalue is deemed good, the scale factor is repeatedly divided by 10 until at least two eigenvalues are deemed good.) Finally, we fit a line using least-squares in log-log space to the data points referred to by the indices, considering only the good eigenvalues. The slope of the fitted line gives the power-law exponent.

### Empirical data

#### Data preparation

We demonstrate GSN on example data taken from the Natural Scenes Dataset (NSD) (Allen et al., 2022). NSD consists of 7T fMRI measurements (1.8-mm resolution) from 8 healthy young adults who each viewed 9,000–10,000 distinct natural scenes up to 3 times each over the course of 30–40 scan sessions. Images were presented for 3 s with 1-s gaps in between images. Participants fixated centrally and performed a long-term continuous recognition task on the images. The fMRI data in NSD come already pre-processed and analyzed using a general linear model as implemented in GLMsingle (Prince et al., 2022). This general linear model produces single-trial beta weights representing the amplitude of the fMRI response on each trial in units of percent signal change. Note that GLMsingle denoises the signal-trial beta weights (i.e. removes some unwanted sources of variance); hence, the analyses in this paper assess the noise that remains after the GLMsingle procedure.

For the purposes of this paper, we took the betas_fithrf version of the single-trial betas in the fsaverage preparation of NSD (the betas_fithrf version reflects a general linear model that accounts for voxel-to-voxel variation in the hemodynamic response function). From the single-trial betas, we extracted responses from several brain regions in the right hemisphere: fusiform face area (FFA-1 subdivision), V1, hV4, and parahippocampal place area (PPA). We use the first region (FFA-1) as the main example; results for the other regions (V1, hV4, PPA) are shown in **S4 Figure**. All regions were functionally localized in each participant, and are supplied with the NSD dataset. We normalized the data by *z*-scoring the responses of each vertex in each session, and then extracted responses for all images that were shown all three times to the participant. (The term ‘vertex’ refers to a point that belongs to a cortical surface representation; for all practical purposes, ‘vertex’ can be treated as synonymous with ‘voxel’ in this paper.) This procedure yielded, for each participant, a set of response measurements with dimensionality *n* vertices × *c* images × 3 trials. As an example of actual numbers, for FFA-1, across participants, the value of *n* ranged from 167 to 1,231 and the value of *c* ranged from 5,445 to 10,000.

#### Application of GSN

We performed GSN on the response measurements from each participant. For the example participant shown in **Figure 6**, GSN was applied to the full dataset as well as data subsets of varying sizes in order to examine the impact of amount of data on estimation quality. This was accomplished by varying the fraction of images used: 1 (10,000 images), 1/4 (2,500 images), 1/16 (625 images), 1/64 (156 images), and 1/256 (39 images). The images in the data subsets were randomly selected and mutually exclusive across subsets. For the full set of participants shown in **Figure 7**, GSN was applied to the full dataset as well as split-halves of the data from each participant. Splitting was performed such that a random half of the images were used for one split and the remaining images were used for the other split.

To aid visual inspection of covariance matrices, we used a particular vertex ordering for the rows and columns of the covariance matrices in **Figure 6**. Specifically, we performed hierarchical clustering (MATLAB’s Statistics Toolbox’s *linkage*.*m*) on trial-averaged responses using a distance metric of one minus correlation and the linkage algorithm of unweighted average distance. This procedure yielded a vertex ordering where similar vertices tend to be close to one another. The same vertex ordering is used for all depicted covariance matrices.

#### Application of PCA

We performed PCA on the results of GSN (‘GSN PCA’). This involved performing PCA separately on the covariance of the signal distribution and on the covariance of the noise distribution. For comparison, we also conducted a naive application of PCA by simply performing PCA on the covariance of the trial-averaged data (‘Standard PCA’).

To compare PCA results across participants, we isolated the set of 515 images that were viewed by all 8 participants 3 times each during the NSD experiment. For each participant, we computed trial-averaged responses for the 515 images and projected these responses onto (i) the principal components associated with the signal distribution in the case of GSN PCA, or (ii) the principal components of the trial-averaged data in the case of Standard PCA. The resulting scores were then compared across participants using the metric of cosine similarity (i.e., the dot product of unit-length-normalized vectors).

One characteristic of PCA is that the sign of each principal component is arbitrary. We performed several sign adjustments to facilitate comparison of PCA results across data splits and participants. First, for every principal component, we flipped the sign of the principal component if necessary to ensure that the mean of the values in the principal component is positive. This incurs no loss of generality and establishes a reasonable starting point for the determination of signs. Second, for corresponding principal components in the split-half analysis for each participant (e.g., PC1 from one half and PC1 from the other half), we flipped the sign of one of the principal components if necessary to ensure that the cosine similarity between the two principal components is non-negative. This flipping procedure ensures that the reliability of results across split halves is not penalized for incidental variation in signs. Third, when comparing scores across participants, we performed a simple iterative algorithm in which scores are sign-flipped if necessary to ensure that the cosine similarity between the scores from a given participant and the average of the scores from the other seven participants is non-negative. This procedure compensates for the sign ambiguity of the principal components derived from each participant.

## Data and code availability statement

The code used in this study is provided at https://osf.io/wkyxn/. The empirical fMRI data used is available at http://naturalscenesdataset.org. The GSN code toolbox is available at https://github.com/cvnlab/GSN/.

## Author Contributions

K.K. developed methods and performed data analysis. H.S. developed methods. J.S.P. and T.G. developed concepts and performed data analysis. G.T. aided in code implementation. K.K., J.S.P., and H.S. wrote the paper. All authors discussed and edited the manuscript.

## Acknowledgements

We thank J. Wilson and B. Pig for comments on the manuscript. This work was supported by NIH grant R01EY034118 (K.K.). Collection of the NSD dataset was supported by NSF IIS-1822683 (K.K.) and NSF IIS-1822929 (T.N.).

---

**S1 Figure.**
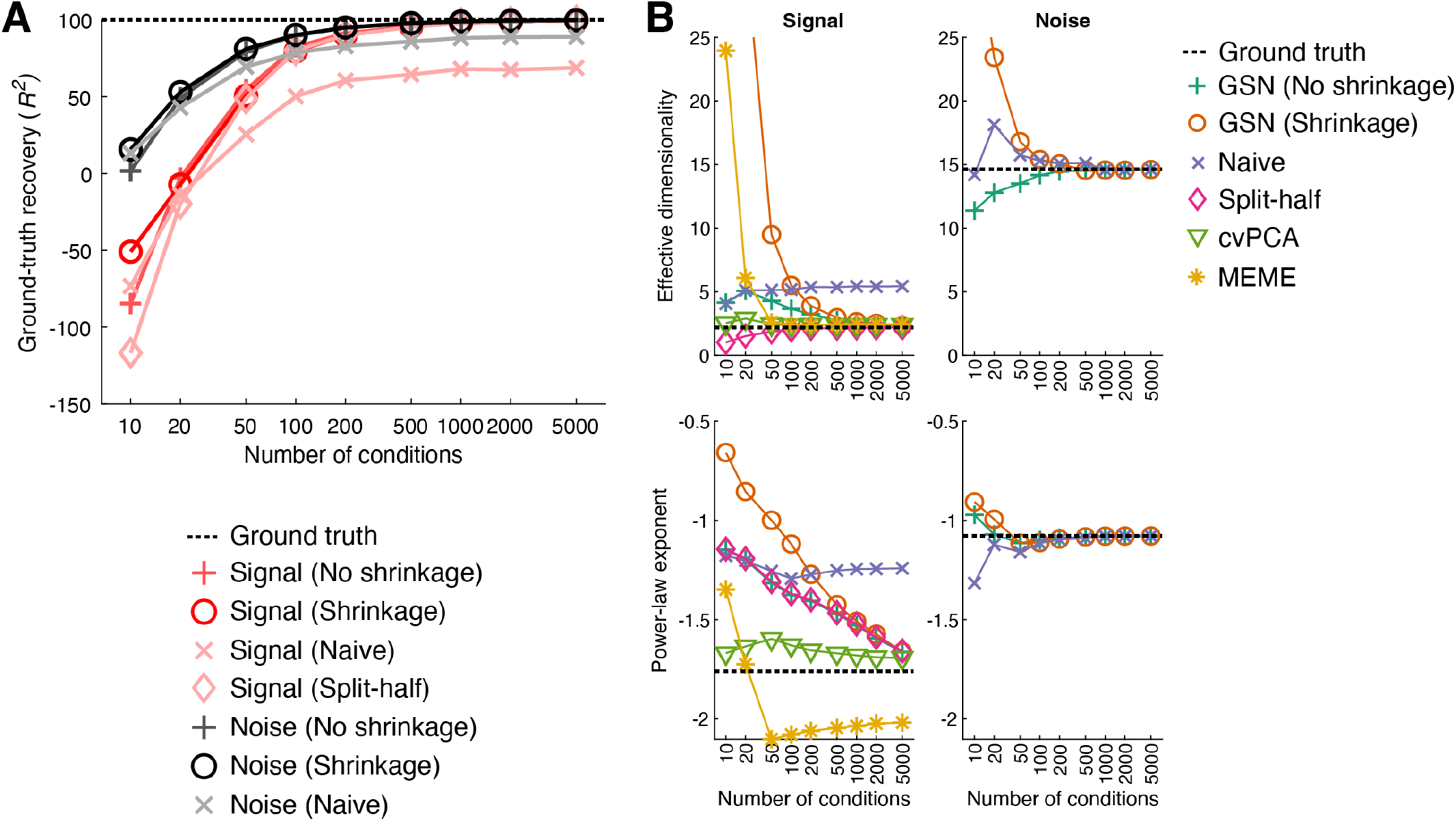
Simulations for empirically derived signal and noise covariance. Here, we show simulation results for a scenario in which we take the ground-truth signal and noise covariance to be the GSN estimates of signal and noise covariance obtained from FFA-1 as illustrated in **Figure 6** (code available at https://osf.io/3yvtg). *A*, Same format as **Figure 4C**. Results are similar to those found in **Figure 4C**. *B*, Same format as **Figure 5**. The results for effective dimensionality (ED) look similar to those found in **Figure 5**. However, the results for power-law exponent look different. Specifically, the methods exhibit poor recovery of signal power-law exponent: each method either takes a very large amount of data to converge towards ground truth or has biases that do not resolve with additional data. One potential explanation is that the ground-truth signal covariance in this scenario is not exactly a line in log-log space (i.e. a power-law function), whereas all of the scenarios shown in **Figure 5** are exactly linear in log-log space. Hence, recovery may be especially difficult to achieve for the current scenario. Arguably, ED is a more appropriate metric for the evaluation of methods here, as it makes minimal assumptions about the structure of the eigenspectrum. Also, note that for sake of consistency with the other simulations, the MEME method was run assuming an unbroken power-law function; in theory, the MEME method could be run assuming a broken power-law function, which might help better match the ground-truth signal eigenspectrum and improve results.

**S2 Figure.**
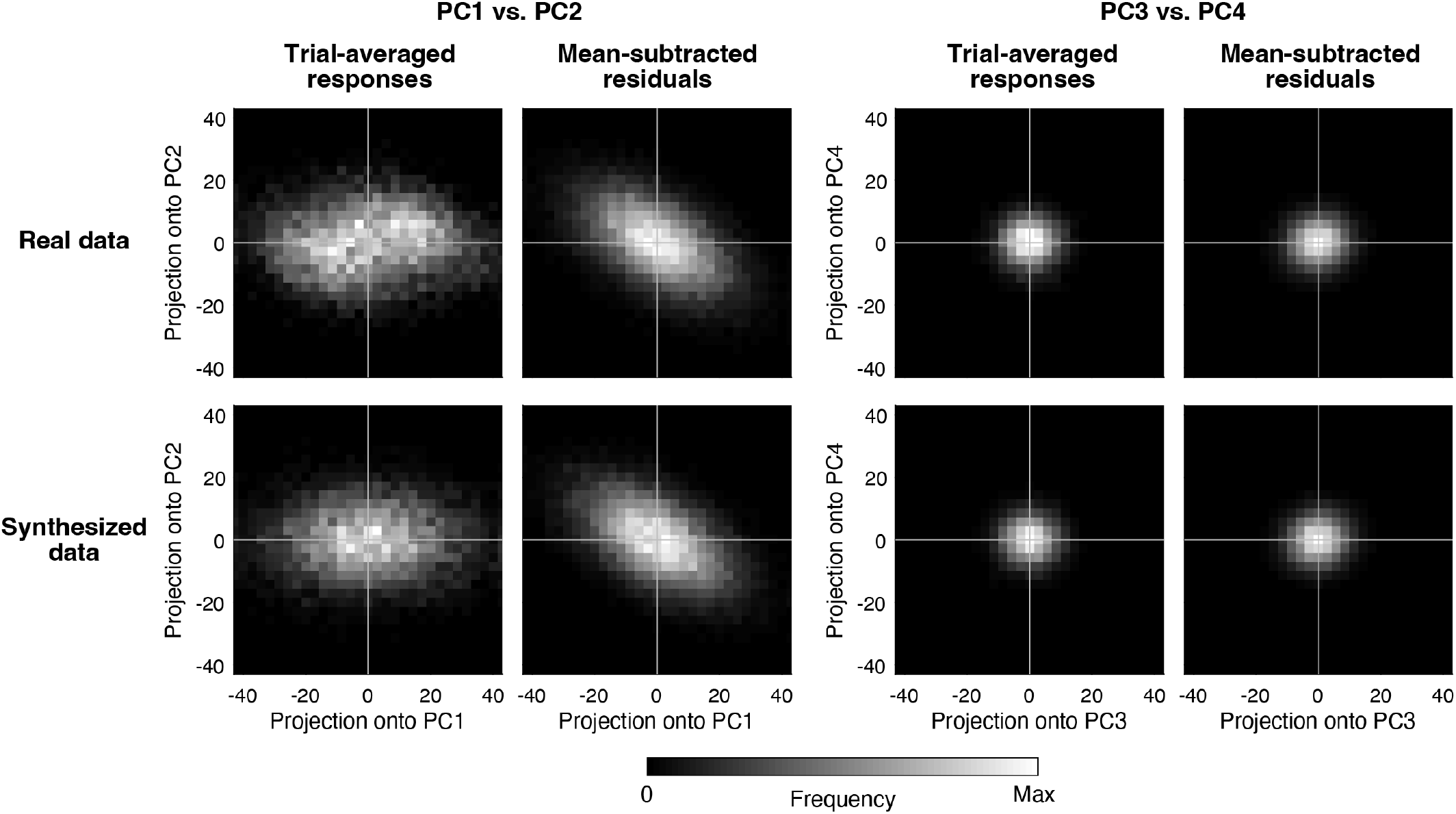
Assessment of data distributions. As an instructive exercise, we take the empirical brain data from FFA-1 illustrated in **Figure 6** and perform an inspection of the signal and noise components of the data (code available at https://osf.io/yxrsp). We inspect two different distributions. One is the distribution of trial-averaged responses. Since trial averaging reduces noise, inspecting trial-averaged responses helps us assess properties of the signal. The second is the distribution of mean-subtracted residuals (in which trial-averaged responses have been removed). This allows us to focus our assessment on properties of the noise. We compute the first several principal components (PCs) of the covariance of the trial-averaged responses and then visualize the two distributions of interest in the low-dimensional space defined by these PCs. We also generate, for comparison, a synthesized dataset based on the parameters of the GSN model as fit to the empirical data. In order to generate responses for this synthesized dataset, we assume that both the signal and noise are Gaussian-distributed. We visualize the synthesized data in exactly the same manner as the real data (including using the same low-dimensional space). Examining the distributions associated with the real data (top row), we see that both the distribution of trial-averaged responses and the distribution of mean-subtracted residuals are Gaussian-like in their shape. We also see that the structure of the mean-subtracted residuals differs from that of the trial-averaged responses (top row, compare first and second images). This indicates that the noise structure is not identical to the signal structure, consistent with the inspections in **Figure 6A**. Next, we compare the distributions associated with the real data (top row) with those obtained from the synthesized data (bottom row). The distributions obtained from the synthesized data look very similar to those from the real data, suggesting that both the signal and the noise in the real data have Gaussian-like distributions and that the generative model learned by GSN accurately characterizes the real data.

**S3 Figure.**
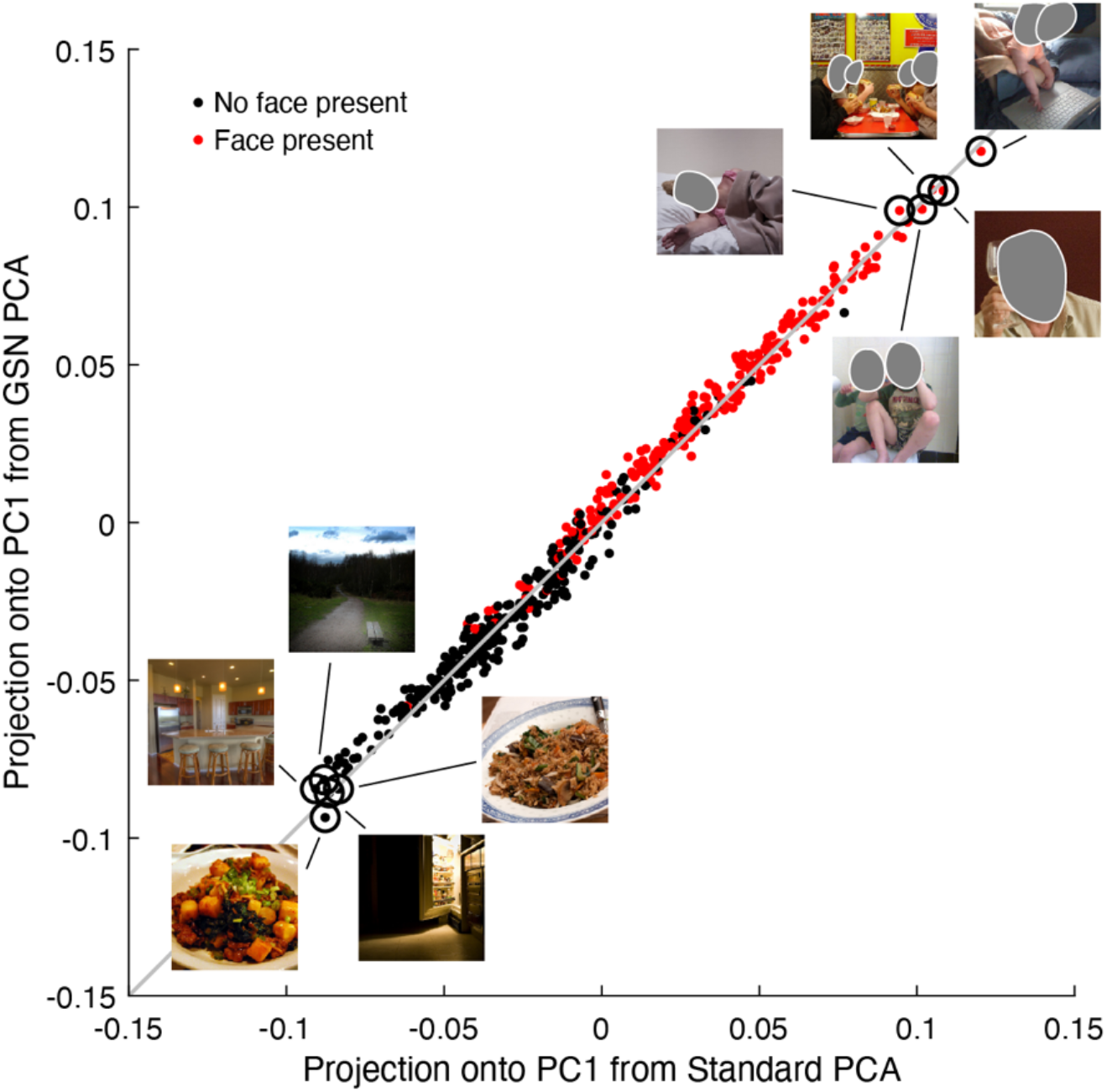
Inspection of stimuli in PCA results. Here we visually inspect stimulus images from the PCA analysis (code available at https://osf.io/f34bc). The projections of responses in FFA-1 to the common 515 images onto PC1 (see **Figure 7C**) were unit-length-normalized, averaged across participants, and then unit-length-normalized again. This figure compares the results obtained using Standard PCA (*x*-axis) against the results obtained using GSN PCA (*y*-axis). Red dots indicate images that were judged by human raters to have at least one prominent face present; black dots indicate all other images. (The human raters were blind to the results in this paper.) The actual images corresponding to the highest five and lowest five projection values (based on the average of the results of the two methods) are shown. The presence of faces appears to be the dominant factor governing the response projections. (Note: Faces have been grayed out due to privacy reasons.)

**S4 Figure.**
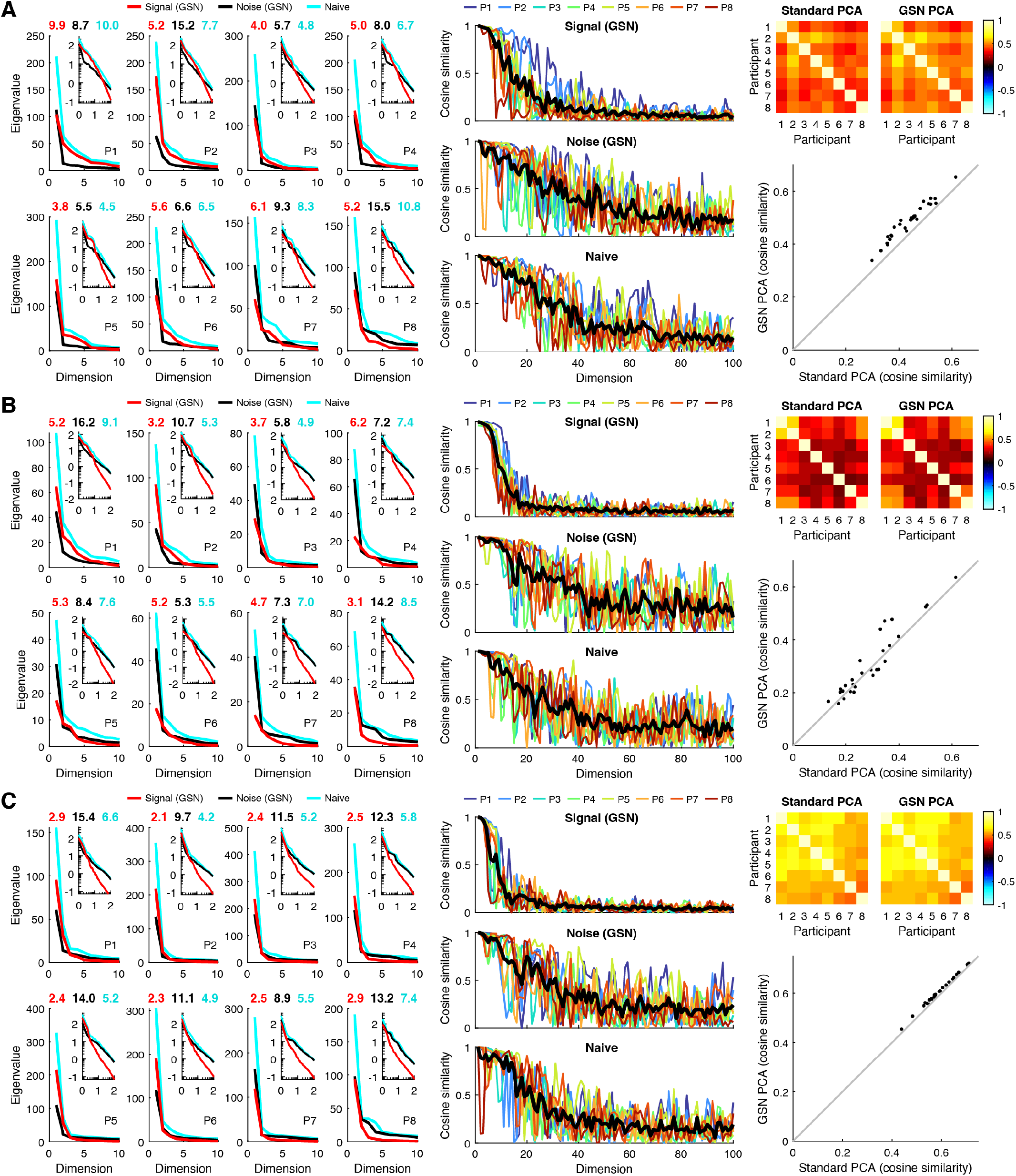
PCA results for additional brain regions. Here we show results of the PCA analysis for additional brain regions (code available at https://osf.io/f34bc). The format is the same as used in **Figure 7**. *A*–*C*, Results for V1, hV4, and PPA, respectively. The main findings observed for FFA-1 in **Figure 7** replicate for these additional regions, including lower dimensionality for the signal compared to the noise, high within-participant reliability of the first several signal PCs and noise PCs, and higher across-participant consistency of trial-averaged response projections onto PC1 for GSN PCA than for standard PCA. Compared to FFA-1, the increase in across-participant consistency is more variable in hV4 and is relatively small (but reliable) in PPA. One possible source of these region-wise differences may be differences in the degree to which signal covariance structure and noise covariance structure are aligned in different brain regions. For example, if noise covariance tends to align with signal covariance, then noise may have less of a corrupting influence on the estimation of signal PCs compared to when noise covariance is orthogonal to signal covariance.

## S5 Appendix: GSN estimation of signal and noise covariance

### Problem setting

As described in the main text, GSN calculates two covariance estimates from the data: 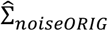 and 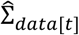. The former is an estimate of the noise covariance based on the trial-to-trial variability around the mean response to each condition (see Step 2). The latter is an estimate of the data covariance based on the data after averaging across *t* trials (see Step 3).

These two covariance estimates reflect unknown covariance matrices Σ_*signal*_ and Σ_*noise*_ such that 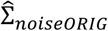 is a noisy version of Σ_*noise*_ based on *c*(*t*−1) samples and 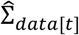 is a noisy version of Σ_*data*[*t*]_ = Σ_*signal*_ + Σ_*noise*_/*t* based on *c*− 1 samples.

We wish to determine estimates 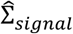 and 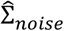 under the constraint that these estimates are positive semi-definite matrices. To do so, we define the following loss that quantifies errors from the data-derived covariances scaled by the number of samples they are based on:

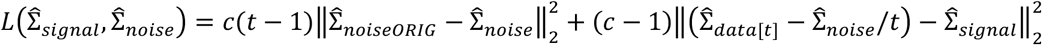

where ‖ ‖_2_ indicates the Frobenius norm. Intuitively, the noise estimate 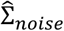 is allowed to deviate to some degree from the data-derived 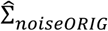, and the signal estimate 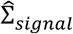 is allowed to deviate to some degree from the subtraction-based estimate of the signal covariance 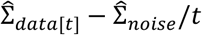.

Notice 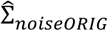 is positive semi-definite, as it is a covariance matrix computed from data. If 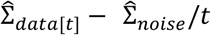 is also positive semi-definite, setting 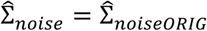 and 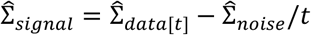 is the optimal solution for the problem (since the loss equals zero). If not, we can solve the optimization problem using the method described below.

### Solution

To solve the optimization problem in the general case, we note that it is a convex optimization problem, as it is a sum of squares and the cone of semi-definite matrices is a convex set (Boyd and Vandenberghe, 2016). Thus, this problem has a single optimum. For solving this problem efficiently, we split the problem into optimizing 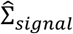 and 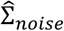 separately, as we can compute an analytic solution for each matrix if the other is fixed. Since each of these separate optimizations is guaranteed to improve the loss, this approach is guaranteed to converge.

#### Lemma: solution pattern

Consider the following problem. Given *B*, find *A* that minimizes 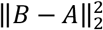 (or equivalently 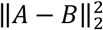) subject to the constraint that *A* is positive semi-definite. We can solve this problem as *A* = PSD(*B*) where PSD() is the method for finding the nearest positive semi-definite matrix described in the main text. We will use this solution pattern in solving the individual optimizations for 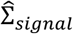 and 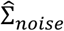.

#### Optimizing 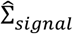

Since 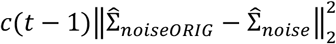 is independent of 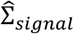, we are left with minimizing

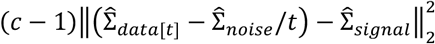

subject to 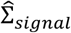 being positive semi-definite. To do this, we use our solution pattern where 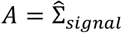 and 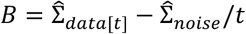.

##### Optimizing 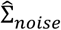

In this case, we apply a quadratic extension to turn the sum of squares into a single one:

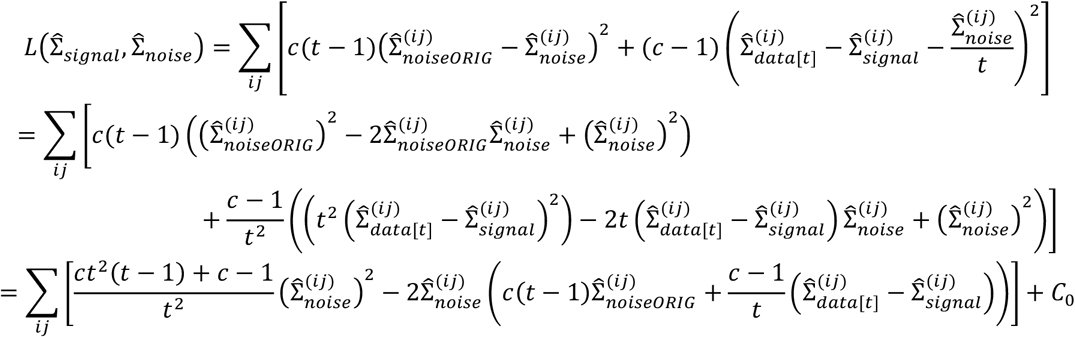

where *C*_0_ is a term that is independent of 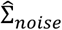. Simplifying, we obtain:

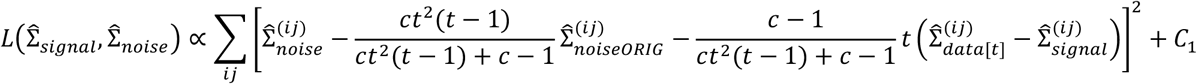

where *C*_1_ is a term that is independent of 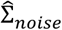. To minimize this loss, we use our solution pattern where

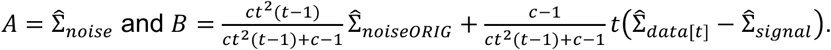

Notice that the calculation of *B* is a weighted average of two possible estimates of the noise covariance. The first estimate, 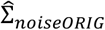, reflects the covariance of mean-subtracted residuals, while the second estimate, 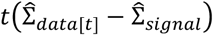, reflects the subtraction of the signal distribution from the data distribution. The weights in the weighted average reflect the amount of data that inform each of the two estimates.

##### Algorithm

The overall algorithm for optimizing signal and noise covariance estimates is described in the main text. Holding 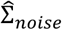 fixed, the algorithm optimizes 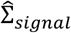 in Step 6.1. Holding 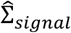 fixed, the algorithm optimizes 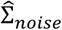 in Step 6.2. This process of biconvex optimization is iterated until convergence.

##### Proof that projection reduces error

We claim in the main text that projection of a given covariance estimate onto the positive semi-definite cone always reduces the error of the estimate. Here we provide a simple proof of this claim.

*Definitions:* For this proof, let Σ be the true *n*-dimensional covariance which lies within the convex cone of positive semi-definite matrices *C* ⊂ ℝ^*n*×*n*^. We assume the original covariance estimate 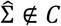.

*Theorem:* Under these conditions, the squared error of the projection onto the positive semi-definite cone 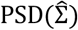 is smaller than the squared error of the original estimate, i.e.:

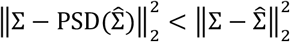

*Proof:* As *C* is convex, there is a tangent plane touching *C* at 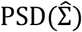 to which the vector from 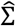 to 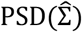 is orthogonal. All points in *C* are on the other side of this tangent plane compared to 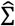. The squared distance from 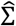 to Σ can be decomposed into the distance orthogonal to the tangent plane and the distance within the tangent plane. The distance within the plane is the same for 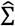 and 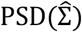, and the distance orthogonal to the plane is smaller for 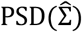. Thus, the total distance for 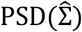 is indeed smaller than the total distance for 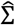. See **Figure S5.1** for a helpful illustration.

**Figure S5.1.**
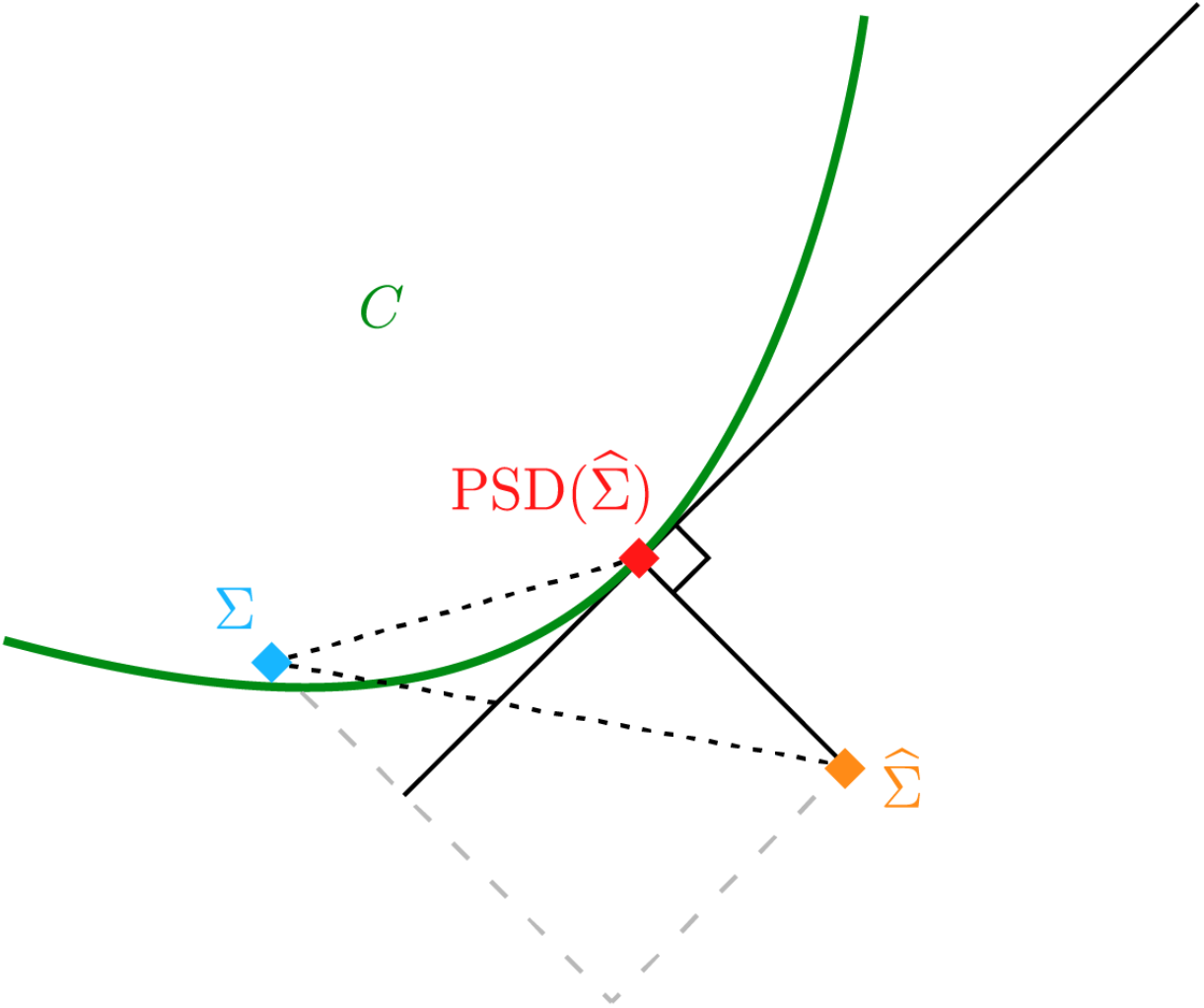
Illustration that projection onto the positive semi-definite cone reduces error.

##### Rationale for squared error

Our estimates are based on minimizing sum of squares, i.e., we minimize the squared difference between our estimates and the data-derived 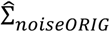 and 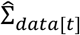. Squared error is a common loss for the estimation of covariance matrices, and in particular, it is the loss optimized by the shrinkage method we employ for covariance estimation. Additionally, squared error is a convex loss function, which guarantees that our fitting procedure converges.

We note that our squared-error loss does not correspond to a log likelihood under some distributional assumption. Rather, it is merely a mathematically convenient way to express the trade-off between the two data-driven covariance estimates 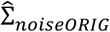 and 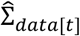. Typical likelihood functions for covariance matrices imply larger variabilities for larger entries in the covariance matrix, but this is not the case for our squared-error loss.

In our squared-error loss, we weight the two errors (one for 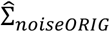, one for 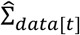) by the relevant degrees of freedom. This is a sensible approach that adapts to the specific numbers of conditions and trials used in a given experiment. We acknowledge that it may be possible to devise a more principled approach for determining the weighting. Nonetheless, note that the relative weighting of the errors does not change fundamental properties of the estimators. For any chosen weighting, 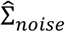 and 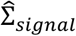 are positive semi-definite and approximate the data-derived covariance estimates.

## S6 Appendix: Shrinkage-based covariance estimation

A core component of GSN is estimation of covariance (this is performed for the estimated noise covariance in Step 2 and the estimated data covariance in Step 3). However, in high-dimensional datasets involving a large number of units but only a limited number of samples (e.g. trials), the standard method of computing sample covariance may yield inaccurate estimates of covariance. To improve accuracy of covariance estimation, GSN incorporates shrinkage (Ledoit and Wolf, 2004; Schäfer and Strimmer, 2005) of off-diagonal elements of covariance matrices towards zero. This reflects the prior that units are generally expected to be uncorrelated. The specific amount of shrinkage is tailored to optimally match the data using a cross-validation procedure in which likelihoods are evaluated on held-out data (see *Methods*).

We tested our shrinkage-based method for covariance estimation. We performed a set of simulations in which we assessed, as a function of the number of samples, how well the shrinkage method recovers a ground-truth covariance, compared to the standard method in which shrinkage is omitted (**Figure S6.1**). In one set of simulations, we used a ground-truth covariance equal to the identity matrix, corresponding to a scenario of uncorrelated units (panel A). In a second set of simulations, we re-used the previous ground-truth covariance but introduced positive correlations (*r* = 0.5) amongst the first five units (panel B). For additional comparison, in a third set of simulations, we used a ground-truth covariance equal to the covariance of a fixed set of random numbers drawn from the standard normal distribution (20 observations, 10 variables) (panel C).

The results show that the shrinkage method works well. In each scenario, the introduction of shrinkage improves ground-truth covariance recovery and this occurs regardless of the number of samples (panels A–C, lower left). Note that the size of the improvement varies across scenarios, with larger improvements when the ground truth is consistent with the prior of uncorrelated units (e.g. panel A) than when this is less the case (e.g. panel C). This makes sense: the cross-validation procedure should, in theory, correctly determine that shrinkage should be applied more strongly in situations where the underlying ground-truth covariance involves uncorrelated variables. Indeed, if we examine cross-validation results across different shrinkage levels, we see that in the scenario of uncorrelated variables, the shrinkage fractions yielding the highest likelihood on held-out data are close to 0, indicating large amounts of shrinkage (panel A, vertical red line in rightmost column), whereas in scenarios of correlated variables, the optimal shrinkage levels are closer to 1, indicating small amounts of shrinkage (panels B–C, vertical red lines in rightmost column).

The simulations also reveal insights into how ground-truth recovery performance varies as a function of the amount of data. As the number of samples increases, the induced shrinkage becomes weaker (compare 5 samples to 100 samples in panel C). This makes sense because at small sample sizes, the unregularized (non-shrunken) covariance is so inaccurate that inducing heavy bias improves the estimate. Furthermore, we see that as the number of samples increases, the difference in results between the shrinkage method and the standard method becomes smaller. Thus, shrinkage provides the most benefit when the amount of available data is small. It is important to keep in mind, however, that the covariance estimates produced by shrinkage are by no means perfect and that they contain bias. This can be seen intuitively by visually comparing the shrinkage-based covariance estimates at low number of samples to the ground-truth covariance. While shrinkage increases the overall similarity of covariance estimates to the underlying ground-truth covariance, it does so at the expense of biasing the magnitudes of off-diagonal elements towards zero. The introduction of bias is not necessarily a problem per se, as it depends on the goals of the researcher.

**Figure S6.1.**
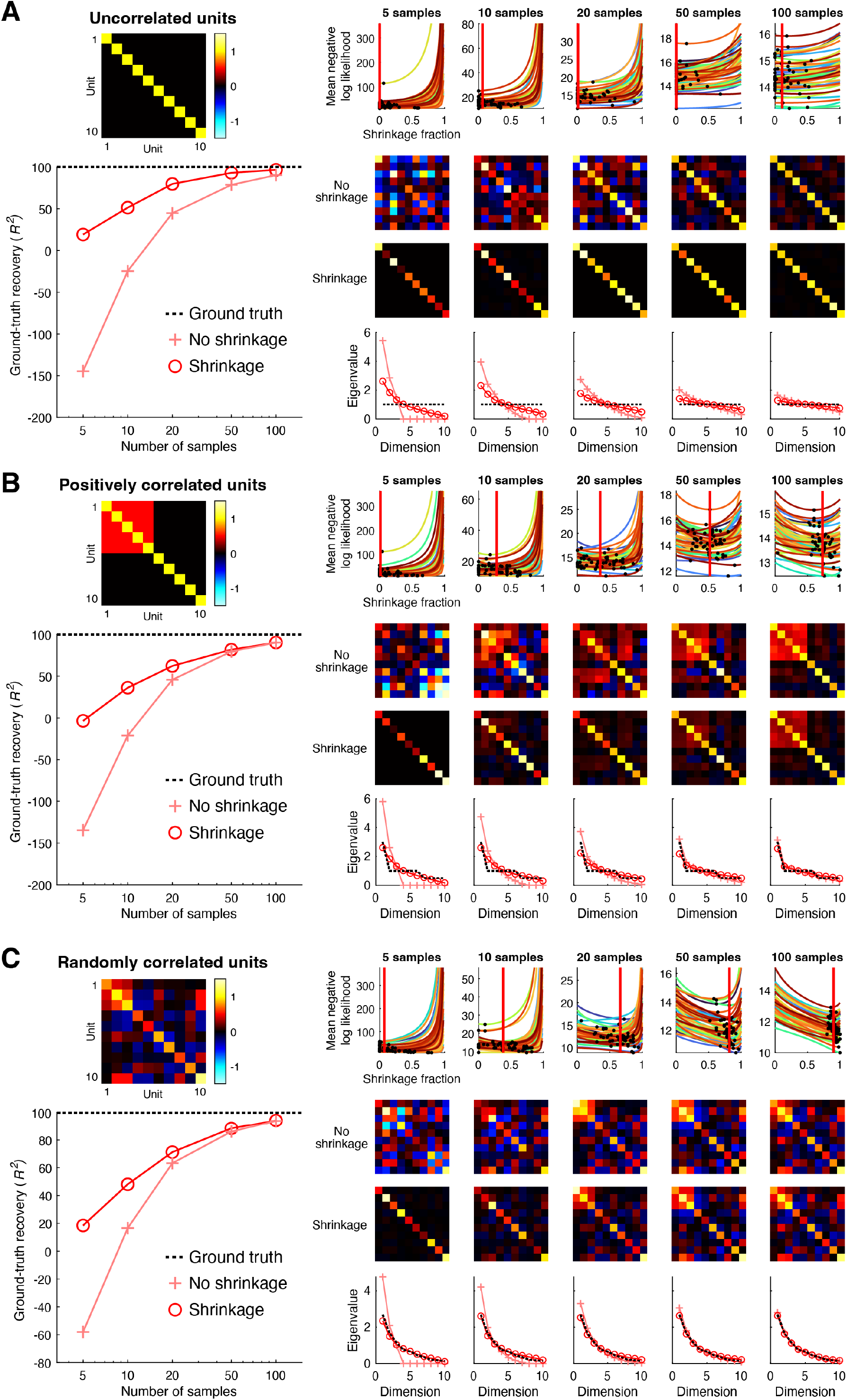
Shrinkage-based covariance estimation. Here we show results of simulations that assess the performance of the shrinkage-based method we use for covariance estimation (code available at https://osf.io/yr3vx). Panels A–C depict three different scenarios. Each scenario involves 10 units whose responses are distributed according to a ground-truth multivariate Gaussian (whose covariance is shown at the upper left). We vary the number of samples (e.g., trials, conditions) drawn from the distribution, performing 50 simulations for each number of samples. In each simulation, we estimate covariance from the samples using two different methods. One method (‘No shrinkage’) is to simply compute the sample covariance with Bessel’s correction. The second method (‘Shrinkage’) involves additionally shrinking the off-diagonal elements of the sample covariance, using cross-validation to determine the optimal shrinkage level. In each panel, the ground-truth covariance is shown at the upper left. Cross-validation results for different numbers of samples are shown at the upper right, where colored lines indicate different simulations, black dots indicate the minimum negative log likelihood achieved, and the vertical red line indicates the median selected shrinkage level across simulations. Below each cross-validation plot, covariance estimates from one simulation are shown (we choose the simulation in which the selected shrinkage level is closest to the median). At the bottom are plots of the eigenspectra (mean across simulations) produced by the two methods (red and pink lines) as well as the ground-truth eigenspectrum (black dotted line). Finally, the ground-truth recovery performance quantified using coefficient of determination (*R*^!^) is shown at the lower left (mean across simulations).

A clear benefit of the bias induced by shrinkage can be seen in the eigenspectra of the covariance estimates (panels A–C, bottom right). Even though the sample covariance provides an unbiased estimate of covariance, it produces biased eigenspectra that are lower in dimensionality than the ground-truth eigenspectra (see steep fall-off of the eigenspectra in the case of 5 samples). In other words, the sample covariance tends to underestimate the true dimensionality of the data. Shrinkage, to an extent, alleviates this issue, as it increases dimensionality (eigenvalues become more spread out) and produces eigenspectra that more closely resemble the ground-truth eigenspectra. These results are consistent with prior results from the literature (see Figure 1 in (Schäfer and Strimmer, 2005)).

